# Interleukin-2 druggability is modulated by global conformational transitions controlled by a helical capping switch

**DOI:** 10.1101/821207

**Authors:** Viviane S. De Paula, Kevin M. Jude, Santrupti Nerli, Caleb R. Glassman, K. Christopher Garcia, Nikolaos G. Sgourakis

**Affiliations:** Department of Chemistry and Biochemistry, University of California Santa Cruz, Santa Cruz, California, USA.; Department of Computer Science, University of California Santa Cruz, Santa Cruz, California, USA.; Howard Hughes Medical Institute, Stanford University School of Medicine, Stanford, California, USA.; Department of Molecular and Cellular Physiology, Stanford University School of Medicine, Stanford, California, USA.; Department of Structural Biology, Stanford University School of Medicine, Stanford, California, USA.

## Abstract

Interleukin-2 (IL-2) is a small α-helical cytokine that regulates immune cell homeostasis through its recruitment to a high-affinity heterotrimeric receptor complex (IL-2Rα/IL-2Rβ/γ_c_). IL-2 has been shown to have therapeutic efficacy for immune diseases by preferentially expanding distinct T-cell compartments, and several T_reg_-biasing anti-IL-2 antibodies have been developed for combination therapies. The conformational plasticity of IL-2 plays an important role in its biological actions by modulating the strength of receptor and drug interactions. Through a NMR analysis of milliseconds-timescale dynamics of free mIL-2 we identify a global transition to an auto-inhibitory conformation controlled by an α-helical capping “switch”. Binding to either an anti-mouse IL-2 mAb or a small molecule inhibitor near the AB loop induces changes in dynamics at the core of the structure, and perturbations of the capping hydrogen-bond network abrogate Ab binding by destabilizing the auto-inhibitory conformation. Our results highlight a paradigm for designing precision therapeutics targeting a continuum of IL-2 conformational states.

## INTRODUCTON

Interleukin-2 (IL-2) is a type I α-helical cytokine that functions as a multi-lineage lymphocyte growth factor^1–2^. IL-2 signals through a high-affinity (10 pM) heterotrimeric receptor complex, consisting of the IL-2Rα (also called CD25), IL-2Rβ (CD122), and γc (CD132) chains, and the intermediate-affinity (1 nM) heterodimeric receptor complex, consisting of the IL-2Rβ and γc chains (Fig. 1a)^1–3^. The IL2-Rα subunit is constitutively expressed in high levels on regulatory T cells (T_reg_) and at lower levels on natural killer (NK) cells and resting effector CD8+ T cells, resulting in differential IL-2 potency between different immune cell compartments^4–5^. From a clinical perspective, both IL-2 agonism and antagonism is of considerable importance and it has been used for more than three decades towards immunotherapies of malignancies.

**Figure 1:**
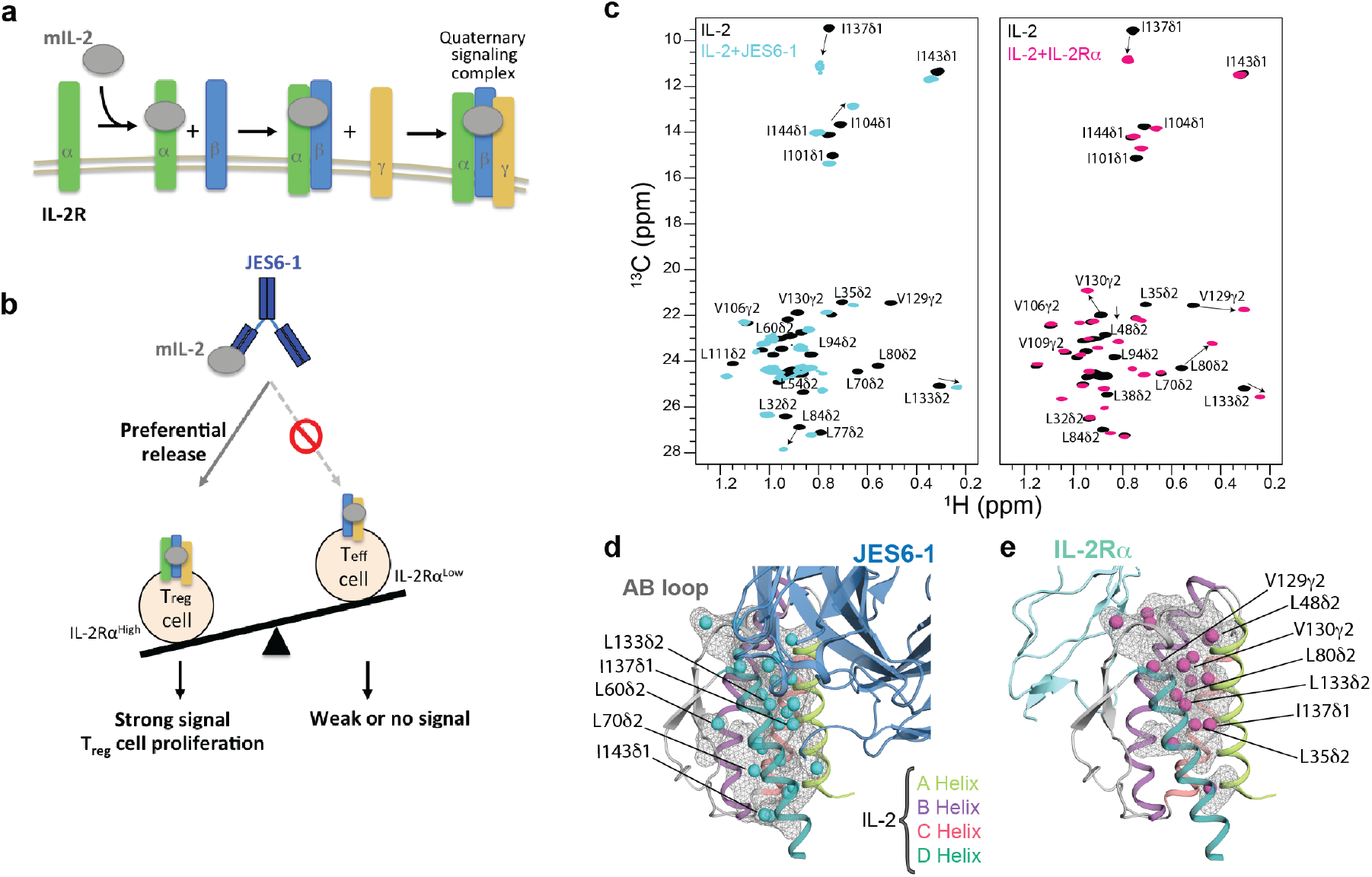
A plastic mIL-2 core structure mediates recognition of its binding partners for immune modulation. **(a)** Schematic of IL-2 cytokine-receptor quaternary complex formation. Assembly of the quaternary complex is thought to occur sequentially, with IL-2 first engaging IL-2Rα with a dissociation constant (*K*_d_) of ∼10^-8^ M, which increases its affinity for the IL-2Rβ subunit, and finally recruiting the γc subunit to lock down the high-affinity quaternary complex (*K*_d_ of ∼10^-11^ M). (**b**) Schematic of the proposed mechanism for mIL-2/JES6-1 immunocomplex-mediated selective proliferation of regulatory T (T_reg_) cells. The JES6-1 Ab (shown as blue scFv) sterically blocks mIL-2 interaction with IL-2Rb and gc subunits, preventing signaling of IL-2Ra^low^ effector cells (right). However, an exchange mechanism between JES6-1 and the IL-2Ra subunit allows a preferential release of mIL-2 for exclusive signaling on IL-2Rα high T_reg_ cells, biasing toward an immunosuppressive response (left). (**c**) Overlay of ^1^H,^13^C-HMQC spectra of selectively labelled mIL-2 at the Iδ_1_-^13^CH_3_, L, V proS methyl positions, recorded in the free (black) or as a stoichiometric complex with JES6-1 scFV (cyan) or IL-2Rα receptor (magenta), acquired at 800 MHz, 25 °C. The arrows highlight major chemical shift effects. Selected methyl assignments have been included. (**d**) Mapping of methyl chemical shift changes on the crystal structure of mIL-2/JES6-1 complex (PDB: 4YQX) and (**e**) on the overlaid IL-2Rα subunit from the homologous hIL-2 quaternary complex structure (PDB: 2B5I). The mIL-2 residues with CSPs > 0.05 ppm are shown with cyan spheres in the cartoon representation of JES6-1-bound mIL-2 (blue) and with magenta spheres in IL-2Rα-bound mIL-2 (green).

The known plasticity of IL-2 has been an important parameter in the generation of small molecules that bind to IL-2 and block signaling^6–7^, engineering ‘super-agonist’ mutant versions of IL-2 with high affinity for IL-2Rβ^8^, and isolation of antibodies that bind to IL-2 resulting in biasing its actions to different immune cell subsets^9–11^. Indeed, many efforts have partially focused in to improve its therapeutic potential by manipulating its ability to selectively target specific cell types^8, 11–16^. In one approach, monoclonal antibodies against IL-2 can alter its properties by binding to a number of distinct conformational epitopes, thereby modifying the interactions of IL-2 with any of the IL-2R subunits and resulting in the proliferation of either T_reg_ or T_eff_ cells^9–10^. For example, wild-type mouse IL-2 (mIL-2) can be administered in complex with an anti-mouse IL-2 monoclonal antibody (JES6-1) and used to preferentially induce T_reg_ cell proliferation^12^. Subsequent work has validated therapeutic applications of the JES6-1 immunocomplex, which promotes graft tolerance^17–18^ and has shown efficacy in preclinical models of diabetes^19–20^. This strategy has been validated for human IL-2 through the discovery of T_reg_-biased antibodies that appear to work by a similar allosteric mechanism^11^. Thus, the conformational states of IL-2 can be selectively induced to achieve cell-type and receptor selectivity, but more detailed insight is needed into the IL-2 conformational transitions.

Focusing on mouse IL-2 as a model system, our team recently determined the X-ray structure of the mIL-2/JES6-1 complex to elucidate the mechanistic basis for its selective stimulation of T_reg_ over effector cells^10^. JES6-1 sterically blocks interactions with the IL-2Rβ and γc subunits leading to a reduction of signaling effects on IL-2Rα^Low^ effector cells. JES6-1 undergoes allosteric exchange with the IL-2Rα subunit, where surface-expressed IL-2Rα displaces the JES6-1 antibody and releases the cytokine to signal through the high-affinity heterotrimeric receptor on IL-2Rα^High^ T_reg_ cells (Fig. 1b). Of particular relevance is the mobility of the AB loop, which has been shown to undergo a large conformational change upon JES6-1 binding. The AB loop is the main binding site for IL-2Rα and crystallographic studies^10^ demonstrated that key residues in the IL-2 AB loop engage the JES6-1 antibody and the IL-2Rα subunit in distinct orientations. Therefore, JES6-1 and IL-2Rα binding are mutually exclusive, leading to bidirectional exchange. Molecular dynamics simulations have suggested a significant conformational rearrangement in the AB and BC loops characterizing the transition from JES6-1-bound to IL-2Rα-bound states^15^. Despite the importance of IL-2 in T cell differentiation and homeostasis, a detailed biophysical characterization of its dynamic properties remains elusive.

In the current work, we use methyl-based Nuclear Magnetic Resonance (NMR) spectroscopy to establish that, in solution, free mIL-2 samples an excited state corresponding to an auto-inhibitory conformation with distinct functional properties. Using a series of complementary NMR experiments we characterize a cooperative transition, which involves a concerted motion of an α-helical capping “switch”, linking the AB loop with the hydrophobic core of the mIL-2 structure through a sequential repacking of core sidechains. Selective disruption of the capping hydrogen-bond network, through a designed mutation, quenches dynamics leading to a 3-order of magnitude loss of affinity for the JES6-1 antibody and, consistently, a loss of immunomodulatory activity *in vitro*. Insights derived from our data highlight the potential of targeting IL-2 excited states using mutagenesis or small-molecule binders, to steer the conformational equilibrium towards states with maximum effects on T_reg_ cells for the treatment of autoimmune diseases, and on T_eff_ cells for immunotherapy of cancer.

## RESULTS

### Long-range effects on mIL-2 core dynamics upon recognition of its binding partners

Due to their hydrophobic character, methyl groups are well-suited NMR probes for monitoring the dynamics of the structural core of proteins^21–22^. Methyl chemical shifts report on sidechain rotameric state, packing and are less influenced by the molecular size of the system under investigation because of their favorable relaxation properties^23^. To gain insight into conformational changes on mIL-2 induced by its binding partners, we prepared selective ^1^H/^13^C MILV (Met, Ile, Leu and Val)-methyl labelled samples on a uniform ^12^C, perdeuterated background. We obtained NMR assignments of mIL-2 backbone and methyl groups using a combination of TROSY-based triple-resonance and 3D out-and-back HMCM[CG]CBCA sidechain transfer experiments. Under the conditions of our NMR experiments mIL-2 is strictly monomeric, as characterized by size exclusion chromatography coupled to multi-angle laser light scattering (SEC-MALS) (**Extended Data Fig. 1**). To obtain stereospecific resonance assignments of methyl groups, and to reduce spectral overlap observed in the free mIL-2 methyl spectrum, which is typical of α-helical proteins, we also prepared a selectively Iδ_1_-^13^C^1^H_3_, proS L,V-^13^C^1^H_3_-labeled protein sample^24^. This approach allowed us to obtain unambiguous sidechain resonance assignments of Met ε_1_, Ile δ_1_, Leu δ_1_/δ_2_ and Val γ_1_/γ_2_ (60 methyl groups in total) (**Extended Data Fig. 1**), distributed throughout the structure, enabling us to comprehensively map the cytokine’s surface and hydrophobic core in the IL-2Rα and JES6-1-bound states. Figure 1c shows ^13^C-^1^H SOFAST HMQCs of I(LV)proS-methyl labeled mIL-2 unbound (black) and as a stoichiometric complex with JES6-1 single-chain Fv (scFv, cyan) or IL-2Rα (magenta). The weighted chemical shift perturbations (CSPs) are presented along the mIL-2 sequence in **Extended Data Fig. 2**. In agreement with previous SPR measurements^10^ under our NMR sample conditions mIL-2 forms tight complexes with either JES6-1 or IL-2Rα. Exchange between the free and bound states is slow on the chemical shift timescale, as indicated by a single set of peaks for each complex, with large CSPs relative to the free state (up to 0.6 p.p.m).

Notably, either receptor or antibody binding lead to measureable conformational changes at the corresponding primary binding sites (for example, L48δ_2_, L50δ_2_ and L54δ_2_ in the AB loop), but also induce long-range effects at remote sites distributed throughout the structure, including L35δ_2_ and L38δ_2_ in the A helix, L73δ_2,_ L80δ_2_, L86δ_2_ in the B helix, I101δ_1_ in the C helix and V129γ_2_, V130γ_2_, L133δ_2_ and I137δ_1_ in the D helix. Mapping the most affected methyl groups on the mIL-2 structure (Fig. 1d, e) highlights a contiguous path connecting the AB loop to the hydrophobic core of the structure. Overall, the methyl CSPs reveal a global response of the mIL-2 structure, indicative of allosteric communication between the primary binding site at the AB loop and core residues of the cytokine (**Extended Data Fig. 2**). Given that the four α-helical bundle core of apo-IL-2 is maintained in the complexed forms with the IL-2Rα receptor or JES6-1 antibody (backbone heavy atom r.m.s.d. of 0.8 Å and 2.3 Å, respectively), our results suggest that binding of either JES6-1 or IL-2Rα at the AB loop region induces a remodeling of core sidechains. Specifically, the observed methyl chemical shift changes can arise from either changes in the lowest-energy rotameric state, or through perturbations in ensembles of rotamers sampled by the free and bound forms.

### Free mIL-2 samples a global transition to an excited-state conformation

Based on our observation that the ^1^H-^15^N transverse relaxation-optimized spectroscopy (TROSY) spectrum of free mIL-2 was of marginal quality (**Extended Data Fig. 1)** with a significant fraction of completely broadened amide resonances (33%), we hypothesized that in solution mIL-2 undergoes exchange between different conformational states. The loss of signal for amide resonances corresponding to residues in the AB loop due to ^15^N line broadening suggests that the loop is switching between multiple backbone conformations with different chemical environments on an intermediate (microseconds to milliseconds / µs-ms) time scale. This hypothesis is consistent with a recent MD simulation-based model in which free IL-2 samples distinct conformations of the AB loop akin to the crystallographically observed structures in complexes with the JES6-1 antibody and IL-2Rα receptor^10, 15^.

To characterize the conformational landscape sampled by free mIL-2 in solution, we performed a series of ^13^C chemical exchange saturation transfer (CEST)^25^ and ^13^C single-quantum methyl Carr-Purcell-Meiboom-Gill (CPMG) relaxation dispersion experiments^26^. Briefly, in CPMG experiments the effective transverse relaxation rate (*R*_2,eff_) is measured as a function of refocusing pulse frequency (νCPMG), which quenches the effects of conformational exchange producing so-called dispersion profiles. Such profiles can be fit to extract the rate of exchange between the ground and excited conformational states, *k*_ex_, the population of the excited state, *p*_E_, and the chemical shift differences between the two exchanging states |Δω| (ppm). In CEST, the intensities of protein resonances are measured in the presence of a weak off-resonance radiofrequency (*B*_1_) field, typically between 5 and 50 Hz, where a series of 2D experiments are acquired with a varying offset of the *B*_1_ field. When the field offset coincides with the resonance of an exchanging minor state, saturation transfer occurs during a fixed period, leading to an attenuation of the resonance of the major state which can be readily detected. This enables the indirect observation of the resonance of the otherwise “invisible”, minor state. While in principle both experiments can be used to quantify conformational exchange processes in biomolecules^27^, CEST is sensitive to timescales ranging from ∼50 to 500 s^-1^, whereas CPMG on the order of 100 to 3000 s^-1^. As a result, CEST experiments are typically recorded at a reduced temperature (4-10 °C), in order to slow the exchange to a measurable rate.

Due to poor quality of the amide spectra of free mIL-2, we turned to methyl spectra as a readout in both CPMG and CEST experiments. Conformational exchange was observed for the resonances of several methyl groups distributed throughout the mIL-2 structure, as illustrated by representative residues on the B helix (L80δ_2_), C helix (I101δ_1_) and D helix (L133δ_1_) (Fig. 2a). The resonances of 20 methyls exhibiting well-defined CPMG dispersion curves (**Extended Data Fig. 3a, Extended Table 1**) were used in a quantitative analysis. The remaining mIL-2 resonances exhibited low S/N dispersion curves (at 18.8 T magnetic field) or high spectral overlap. Dispersion curves were fitted globally using a two-site exchange model, yielding a *k*_ex_ of 1,241 ± 72 s^-1^ and an excited state population of 6.0 ± 0.4%. Additionally, ^13^C-CEST profiles for the same residues (L80δ_2_, I101δ_1_ and L133δ_1_) revealed significant dips in the ^13^C dimension, characteristic of an excited state (Fig. 2b **and Extended Data Fig. 4**). To quantify the extent of structural adaptations upon formation of the excited state for each methyl site participating in the exchange process, we extracted values of the corresponding ^13^C chemical shift changes, |Δω|, from independent fits of the CPMG relaxation dispersion and CEST data sets. We observe good correlation between the resulting |Δω| values, suggesting that the two methods are likely reporting on a similar exchange process, with the caveat that the CEST data were recorded at a lower temperature (Fig. 2c).

**Figure 2:**
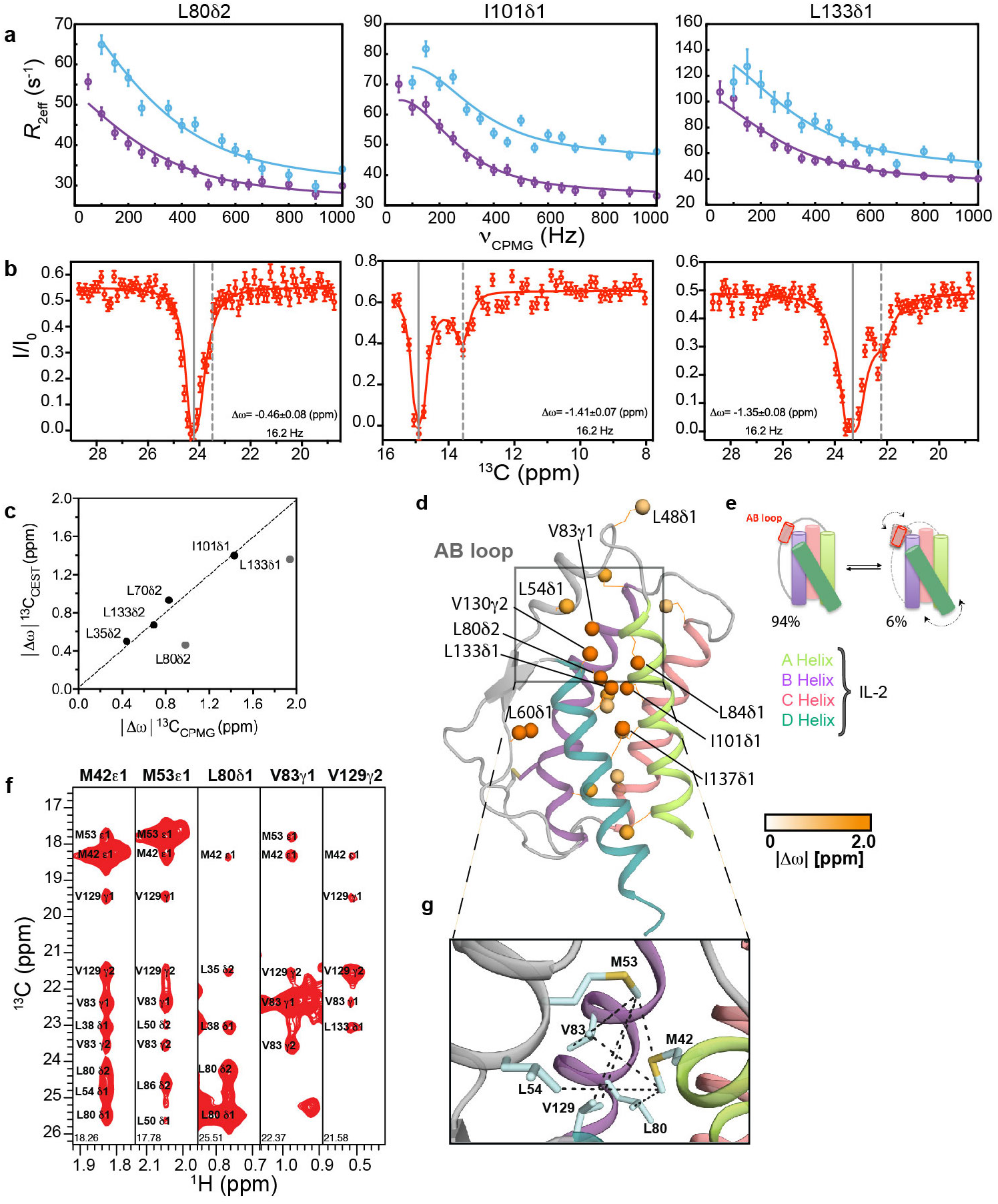
Solution dynamics and ground-state structure of free mIL-2 by methyl NMR. (**a**) methyl-selective ^13^C single quantum CPMG relaxation dispersion profiles carried out at two magnetic fields (600 MHz, purple; 800 MHz, blue) are shown for selected residues in the B helix (L80δδ_2_), C helix (I101δ_1_) and D helix (L133δ_1_) of WT free mIL-2. Experimental data are shown as small circles in all panels, with errors estimated from the S/N in the raw experimental data. The best-fit lines are shown for a global analysis of 20 methyls with non-flat relaxation dispersion profiles using a two-site conformational exchange model. (**b**) ^13^C-CEST profiles carried out at 800 MHz for L80δ_2_, I101δ_1_ and L133δ_1_. Plots of normalized intensity of the ground state resonance relative to a control experiment where the saturating field is highly off-resonance (12 kHz), as a function of the position of the perturbing B_1_ field (with a measured strength of 16.2 Hz). Uncertainties in I/I_0_ are determined from the root-mean-square deviation in the baseline of the raw CEST profiles where no intensity dips are present (typically, n > 30). Global fits of the CEST data to a two-site model of chemical exchange are shown as solid red lines. The resonances of the major state (gray solid lines) and fitted minor state (grey dashed lines) are indicated, with the resulting chemical shift difference shown in each plot. (**c**) Linear correlation plot of chemical shifts of the excited state of WT mIL-2 obtained from ^13^C CPMG (x-axis) and ^13^C CEST (y-axis). CPMG and CEST experiments were recorded at 25 °C and 10 °C, respectively. (**d**) 20 Methyl probes undergoing chemical exchange by CPMG are shown as spheres on the model structure of mIL-2 and colored orange according to the magnitude of fitted |Δω| values. (**e**) Cartoon representation of mIL-2 illustrating the interconversion between a ground state, which is 94% populated, and an invisible, excited state, which is ∼6% populated. The doted arrows represent the coupled conformational transition of the AB loop and the hydrophobic core of mIL-2. (**f**) ^13^C_M_-^1^H_M_ strips from a 3D C_M_-C_M_H_M_ SOFAST NOESY experiment taken at the ^13^C_M_ coordinates of stereospecifically assigned methyl resonances noted on each panel, showing NOE cross-peaks between the methyl resonances of residues on the AB Loop. (**g**) Close-up view of the AB loop region from a *Rosetta* homology-based model of free mIL-2 (using PDB ID 1M47 as a template) showing the network of observed NOEs (black dotted lines), corresponding to the major (ground-state) solution conformation. The pattern of NOEs is consistent with a “closed” conformation of free mIL-2, with where the AB loop is well-packed against the hydrophobic core of the structure.

Methyl groups exhibiting CPMG dispersion curves are plotted on the homology-based model of free mIL-2 in Fig. 2d, and color-coded according to the magnitude of the fitted |Δω| values, which reports on differences in the local magnetic environment between the major and minor conformations. Large |Δω| values were observed for methyls at the AB loop (L48δ_1_, L54δ_1_), in addition to the C-terminus of the B helix facing towards the loop (L80δ_2_, V83γ_1_, L84δ_1_) and throughout the hydrophobic core of the structure (I101δ_1_, V130γ_2_, L133δ_1_ and I137δ_1_). Thus, our CPMG data suggest that free mIL-2 samples a global, cooperative transition to an excited state, which involves a conformational “switch” of the AB loop, coupled to a cooperative repacking of core sidechains (Fig. 2e). Notably, a pronounced structural change relative to the free form can be observed in the X-ray structure of JES6-1-bound mIL-2^10^, in which the AB loop (residues Y45-T55) undergoes a ∼38° rigid-body rotation to adopt a conformation that is optimal for interactions with the antibody surface. Taken together, conformational plasticity of the AB loop is a key component of the solution dynamics of free mIL-2, relevant for the formation of a high-affinity immunomodulatory complex.

To characterize the major conformation sampled by the AB loop in solution, we analyzed methyl NOE intensities recorded in a 3D C_M_-C_M_H_M_ SOFAST NOESY experiment (Fig. 2f), relative to the corresponding distances observed in *i*) a model of free mIL-2 built using the human IL-2 apo-structure as a template, (PDB ID 1M47) *ii*) the co-crystal structure of mIL-2 in complex with the JES6-1 Ab (PDB ID 4YQX) and *iii*) a model of mIL-2 build using the human IL-2/IL-2Rα receptor complex structure as a template (PDB ID 2B5I). Consistently with the models of mIL-2 in the apo and IL-2Rα receptor-bound states, we find that the network of observed NOEs connecting the methyl groups of M42, M53, L54, L80, V83 and V129 located at the vicinity of the AB loop, (Fig. 2g**, black dotted lines**) is consistent with a “closed”, well-packed conformation where the sidechains of M53 and L54 are making hydrophobic contacts with residues at the hydrophobic core defined by the amphipathic A and B α-helices. This is evident from the observation of several unambiguously assigned short-range (5 Å upper limit) NOEs between the corresponding methyl groups (Fig. 2d, f and g). On the contrary, in the antibody-bound mIL-2 structure the AB loop is found in an “open” conformation, with the methyl groups of M53 and L54 oriented towards the solvent at distances from the core methyls (13 Å), well beyond the NOE detection limit of 10 Å (Fig. 2g **and Extended Table 2**). Thus, our NOE data provide strong evidence that the major state of free mIL-2 has a closed AB loop conformation, similar to the X-ray structure of human IL-2 in either the free form or in the complex with IL-2Rα.

### A conserved hydrogen bond network stabilizes the excited mIL-2 conformation

We next sought to characterize the structural features of the excited-state conformation identified by our CPMG and CEST experiments. Excited states play important roles in protein function, including catalysis^28–30^ and complex formation via conformational selection or induced fit-type mechanisms^31–32^. However, their *de novo* structure modeling is challenged by the inherently low-resolution nature of the NMR chemical shifts. A qualitative comparison of the placement of residues undergoing conformational exchange in free mIL-2 (Fig. 2d) with those perturbed upon formation of the IL-2Rα- and JES6-1-bound states (Fig. 1d, e) showed a significant overlap, both for residues that are within the AB loop and other binding surfaces (L80δ_2_ and L86δ_2_) and also at the core of the structure (L133δ_2_ and I137δ_1_), suggesting that the same sites that participate in the formation of the excited state also undergo structural changes upon formation of the mIL-2 tertiary complexes.

Our NOE-based analysis has shown that the backbone conformation of the AB loop in the major, “closed” state of mIL-2 is inconsistent with the JES6-1-bound structure. Close inspection of the structure reveals that, in the JES6-1 complex, the AB loop instead participates in a network of C-capping interactions with the A helix (Fig. 3a,d). Specifically, the sidechain guanidino group of R52 forms multiple hydrogen bonds with the backbone carbonyl oxygens of M42, Y45 located at the C-terminal end of A helix, and with R46, L48 at the N-terminal part of the AB loop (Fig. 3b,c). Conversely, in the free structure of the homologous human IL-2 (1M47 - termed “uncapped” state), the R52 sidechain is solvent-exposed, and M53 is buried at the hydrophobic interface between the A and B α-helices. The “closed/uncapped” conformation is optimal for interactions with co-receptors, as exemplified by the IL-2Rα complex where R52 forms a salt bridge with a conserved aspartic acid on the receptor surface (PDB 2B5I). JES6-1 binding therefore induces a transition from a “closed” to “open” AB loop conformation, which exposes the hydrophobic sidechains of L50, M53 and L54 towards the surface of the molecule, while the sidechain of R52 becomes buried. In this “capped” state, the loss of hydrophobic packing contacts is compensated by the formation of multiple C-capping hydrogen bonds with the A helix, and the backbone of the loop itself (Fig. 3a,c).

**Figure 3:**
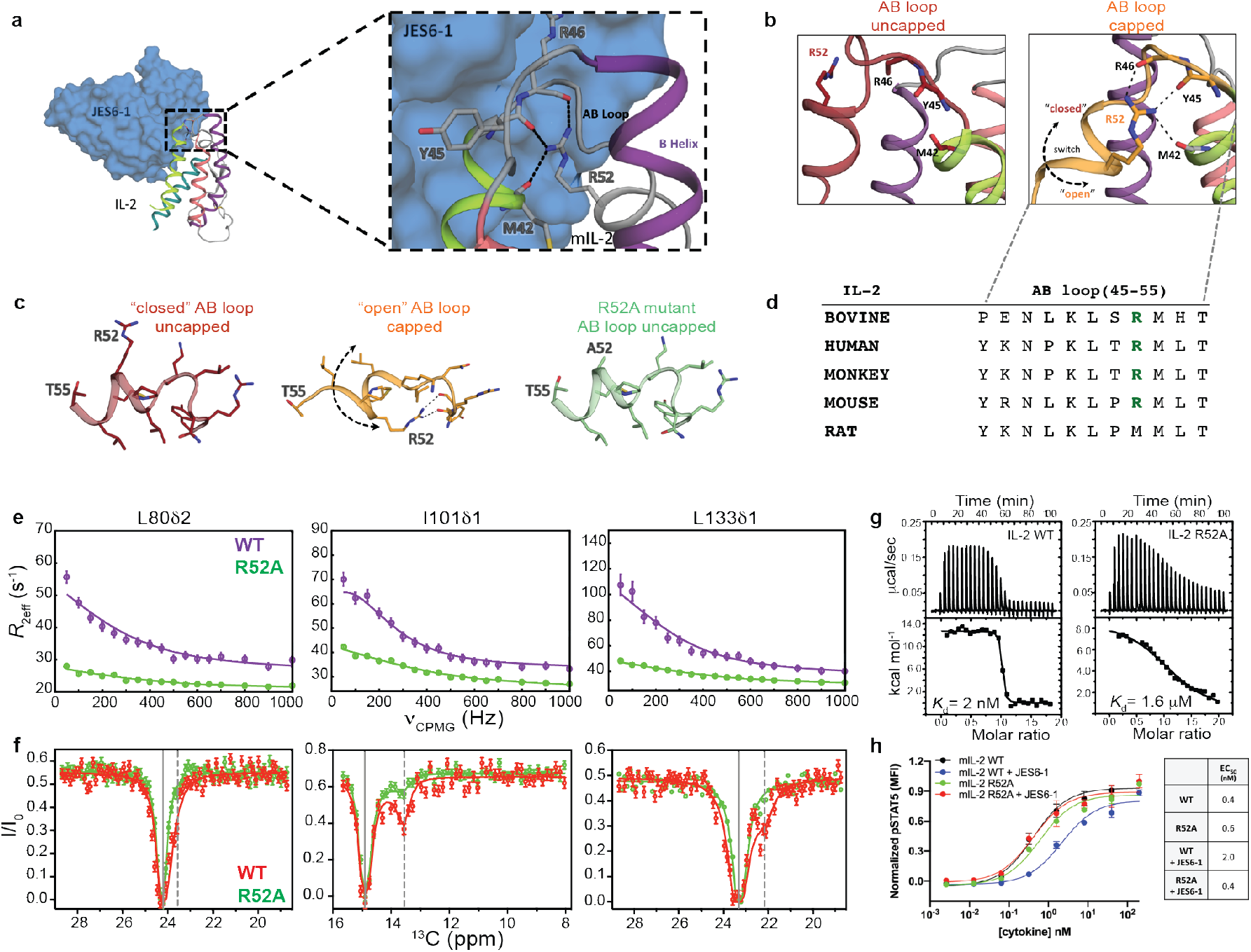
Effects of a conserved helical capping motif on IL-2 dynamics and function. (**a**) Crystal structure of mIL-2 (cartoon) bound to JES6-1 (blue surface) with a detailed view of the mIL-2/JES6-1 interface on the AB loop and B helix of the cytokine (PDB: 4 YQX). (**b**) The AB loop in mIL-2 adopts either an uncapped “closed” conformation (apo state) or a capped “open” conformation (JES6-1-bound state). The open conformation is stabilized by an intramolecular C-capping interaction between the sidechain of R52 and the backbone of M42-Y45-R46. Black dashes indicate hydrogen bonds. The R52 sidechain does not contribute to intramolecular stabilizing interactions in the closed conformation. Black dotted arrow shows the direction of loop movement between the closed and open states. (**c**) Close-up view of the AB loop in the closed and open conformations. The R52A mutation eliminates the C-capping interactions. (**d**) Sequence conservation pattern in the AB loop region in various IL-2 species. (**e**) Comparison of ^13^C-CPMG relaxation dispersion profiles for L80δ_2_, I101δ_1_ and L133δ_1_ of WT mIL-2 (purple) and R52A (green), at 600 MHz and (**f**) ^13^C-CEST profiles for WT mIL-2 (red) and R52A (green). Conformational exchange profiles throughput the mIL-2 structure are quenched by the R52A mutation. Experimental errors in CPMG and CEST data are determined as described in Figure 1. (**g**) Thermodynamic fingerprints of the interactions between WT and R52A mIL-2 with JES6-1. Isothermal titration calorimetry thermographs and curve fits for titrations. The R52A mutation leads to a 3-order of magnitude reduction in binding affinity. To minimize enthalpy of solvation effects, all experiments were performed in 20 mM phosphate buffer, pH 7.2, 150 mM NaCl. (**h**) STAT5 phosphorylation response to WT and R52A mIL-2 in the free state or JES6-1 immunocomplexes treatment in IL-2Rα^+^ mouse CTLL-2 cells. R52A shows attenuated immunomodulation by the JES6-1 Ab, while signaling via IL-2Rα^+^ is at similar levels to WT.

According to our NOE-based analysis, the placement of M53 at the hydrophobic interface between the AB loop and the core of the structure promotes the “closed” loop conformation, which is the major solution state. We sought to determine whether the excited state sampled by free mIL-2 encompasses the C-capping features of the “open” state. If the dynamics observed by NMR correspond to a concerted process, such as an open-to-closed transition of the AB loop, then mutations which destabilize the open state should alter the observed exchange parameters in a uniform manner. To perturb the “open” state conformation, we used the mutant R52A which eliminates all C-capping interactions (Fig. 3c **and Extended Data Fig. 5**) and compared the resulting CPMG relaxation dispersion and CEST data performed under identical conditions to our established results for the WT. Remarkably, Ala mutation of R52 suppressed the CPMG relaxation dispersion profiles of all methyl probes that were undergoing conformational exchange in the WT form (Fig 3e,f **and Extended Data Fig. 3b**). Data fitting showed that, in the R52A mutant, the exchange process was 2-fold faster relative to the WT with a *k*_ex_ of 2,692 ± 80 s^-1^ and had a 3-fold reduced excited state population of 2.0 ± 0.1%. Consistently, in CEST profiles recorded using the R52A mutant the second dip was barely observed (I101δ_1_) or notably absent (L70δ_2,_ I143δ_1_, L133δ_1_, L133δ_2_) (Fig. 3e,f **and Extended Data Fig. 4**), indicating that the population of the excited state (“closed” state) was below the detection limits of the experiment. The R52A mutation quenched µs-ms relaxation not only at the sidechain methyl but also the backbone amide groups, alleviating resonance broadening and leading to a significant improvement in the quality of the ^1^H-^15^N TROSY spectrum (**Extended Data Fig. 5a)**. These observations imply that while at least two conformational exchange processes are at play on the µs-ms time scale (AB loop switching and a transient twisting or ‘‘breathing’’ of the α-helices), the conserved C-capping sequence motif at the AB loop is critical for the formation of the excited state, which is directly coupled to changes at the core of the structure.

To determine how destabilization of the excited state affects mIL-2 binding to JES6-1, we used isothermal titration calorimetry (ITC) and measured the binding free energy (ΔG), and its enthalpic (ΔH) and entropic (-T*ΔS) components both for WT and R52A mIL-2 (Fig. 3g,h). Consistently with previous SPR measurements^10^ and our observation of a tight complex in slow-exchange by NMR, WT mIL-2 binds JES6-1 with a high affinity of 2 nM. The interaction is largely entropy-driven, with a positive net enthalpy change of 12.7 kcal mol^-1^ and a favorable entropic contribution of −24.3 kcal mol^-1^ at 293 K (Fig. 3h**, Extended Table 3**). Notably, R52A binds to JES6-1 with 3 orders of magnitude lower affinity than WT mIL-2, due to less favorable binding entropy (Fig. 3g,h). The weaker association cannot be due to a loss of interface contacts or buried surface area, given that R52 does not participate in direct interactions with JES6-1 in the X-ray structure of the complex (Fig. 3a). Thus, our ITC results show that perturbations of the capping hydrogen-bond network abrogates JES6-1 binding by destabilizing the open conformation, consistently with our NMR measurements showing a shift of the conformational equilibrium towards the closed state.

To determine the functional activity of the R52A mutant, we probed JES6-1 antibody effects on mIL-2-induced signaling in IL-2Rα^+^ subpopulations of mouse CTLL-2 cells as a surrogate for its effects on IL-2Rα^hi^ immune cell subsets (Fig. 3h). Consistent with our NMR and ITC results, JES6-1 showed a 5-fold reduced WT mIL-2-mediated STAT5 activation. However, STAT5 signaling potency by R52A was independent of JES6-1, as demonstrated by the comparable EC_50_ values, suggesting that the weaker affinity to JES6-1 was primarily responsible for the behavior of the R52A mutant (Fig. 3g).

### Allosteric communication in mIL-2 through a remodeling of sidechain rotamers

Our methyl-based NMR probes revealed a global response of the mIL-2 structure upon binding to JES6-1, alongside the crystallographically observed conformational changes of the AB loop (Fig. 1c, d). Consistently, our NMR relaxation measurements suggest a coupling of dynamic motions between the loop and core methyls. We sought to identify a plausible mechanism linking AB loop movement to changes in core sidechain packing during transitions between “closed/uncapped” and “open/capped” conformations. To enumerate all possible sidechain rotamers that can be adopted by each residue, we performed a global analysis of compatible rotamer pairs using a satisfiability-based approach in *Rosetta* (see Methods, **Extended Data Fig. 7**) and mapped our results on the free and JES6-1-bound mIL-2 structures (Fig. 4**, Extended Data Fig. 8**). Using the backbone conformations of the “open” and “closed” states as inputs, our analysis highlights differences in rotamer sets that can be accessed by the free (closed) and JES6-1 bound (open) states. We identified a large set of residues (L28, L39, M42, L48, M53, F58, F132, L133, W136 and F139) (Fig. 4a, b) spanning the AB loop, A Helix and D Helix, and part of the hydrophobic core. For these residues the space of rotamers was significantly different between the closed and open states, indicating a plausible remodeling of packing interactions. Specifically, a 10-residue segment (V129-F139) forming the hydrophobic face of the amphipathic D helix exhibits expansions and contractions in allowed rotamer sets as IL-2 transitions between the two states (**Extended Data Fig. 7**). A concerted motion of the D helix upon binding of IL-2Rα has been highlighted as having a functional role in the formation of the high-affinity heterotrimeric signaling complex (Wang et al., 2005). These results are consistent with our observed NMR chemical shift mapping of the IL-2Rα and Ab-bound states (Fig. 1c, d) suggesting that the conformational transition between the free and complexed states can lead to a redistribution of the rotameric states at the hydrophobic core, albeit limited by known ambiguities in interpreting methyl chemical shifts. A plausible allosteric communication network starts at the AB loop on one end of the structure, traverses through the inner core of mIL-2 and ends at the N-terminus of the A and D helices adjacent to the binding site of the IL-2Rβ receptor (Fig. 4c). The hydrophobic core residues exhibiting differences in rotamer sets also include sites with significant *R*_ex_ values in our NMR CPMG data, indicating the presence of dynamics at the µs-ms timescale (Fig. 4c, orange circles). Taken together, our results highlight a plausible allosteric communication network in the IL-2 structure mediated via sequential remodeling of sidechain packing interactions.

**Figure 4:**
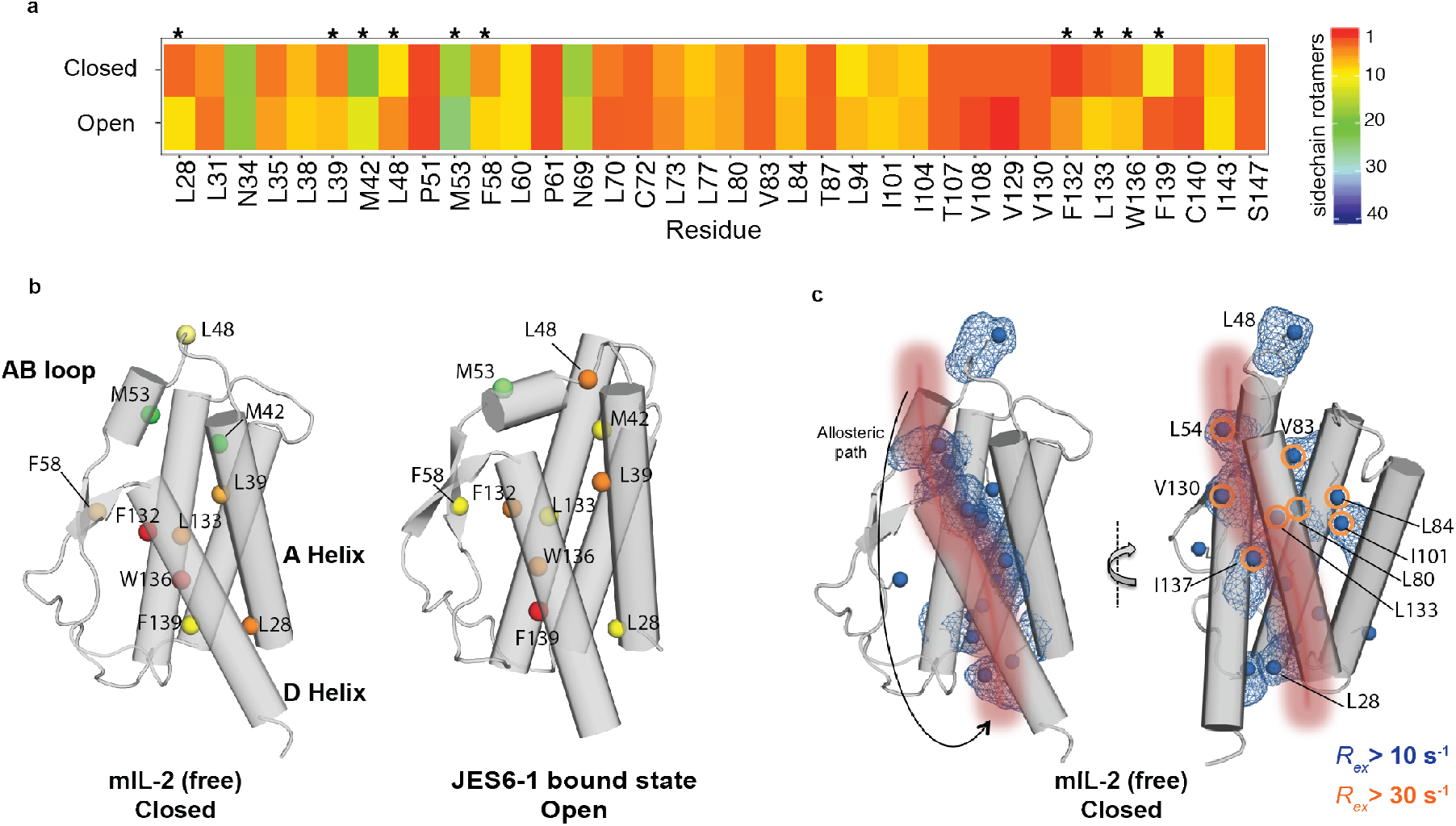
Sampling of different ensembles of sidechain rotamers in the “open” and “closed” mIL-2 states. **(a)** Heat map showing the number of allowed sidechain rotamers for buried residues in the “closed” and “open”-state structures. Buried residues were computed using a 10 Å^2^ solvent accessible surface area threshold. Residues that show a significant difference (> 3) in the number of allowed rotamers between closed and open states are highlighted with stars (*). (**b**) Closed/free and open/bound structures showing residues that exhibit significant difference in sidechain rotamer sets, with the total number of allowed rotamers colored as in panel a. (**c**) Illustration of a putative allosteric communication network linking the AB loop conformation to the core structure. A sequential path demarcated by residues undergoing sidechain remodeling as mIL-2 transitions between the two states is shown with a red patch onto the mIL-2 structure of the closed state, used as a reference. The methyl groups showing significant exchange (*R*_ex_) contributions in our CPMG experiments indicating dynamics are highlighted with blue surfaces (*R*_ex_ > 10 s^-1^) or orange circles (*R*_ex_ > 30 s^-1^).

### Skewing the dynamic landscape of mIL-2 by ligand binding

The R52A mutation characterized here destabilizes the open mIL-2 conformation by perturbing the C-capping hydrogen-bond network between the AB loop and B helix, leading to quenching of conformational exchange throughout the core of the structure. We hypothesized that a small molecule binding preferentially to the closed AB loop conformation would impact the dynamic landscape of mIL-2 in a similar manner. We used a known compound targeting hIL-2 (Ro 26-4550), to compete with IL-2Rα binding^7, 33^. The co-crystal structure (PDB 1M48) shows that Ro 26-4550 is nestled in a hydrophobic pocket at the interface between the AB loop and B helix, with the hIL-2 sidechains of M39, V69, and L72 (mIL-2 residues M53, V83 and L86) packing against the terminal aromatic ring^7^. We performed an NMR titration of increasing molar ratios of Ro 26-4550 on ILV-methyl labeled WT mIL-2 using ^1^H-^13^C HMQC spectra as a readout (Fig. 5a). Under our NMR conditions, we observed the formation of a moderate-affinity (µM range K_d_) complex in fast exchange with the free mIL-2 form, where residues in close proximity to the AB loop (L48δ_1_, L54δ_1_ and L60δ_1_) experience the largest effects (**Extended Data Fig. 6**). However, CSPs extend beyond the AB loop, to sites in the hydrophobic core (L80δ_1_, V129γ_2_, V130γ_2_, L133δ_2_ and I137δ_1_) at distances >12 Å from the inhibitor binding site in the co-crystal structure (Fig. 5c). A quantitative NMR line shape analysis of the resonances of V129γ_2_ and L133δ_2_ yielded an equilibrium dissociation constant of 39.4 ± 5.5 µM (Fig. 5b), suggesting a 2-state cooperative transition impacting all sites in the ligand-bound form. To further examine whether Ro 26-4550 binding affects the µs-ms dynamics of mIL-2, we repeated ^13^C-CPMG relaxation dispersion experiments under saturation binding conditions under identical protein concentration and NMR sample conditions, and compared with our established data recorded for the free form (Fig. 5d). Our results indicate that Ro 26-4550 binding strongly dampens relaxation of methyl groups that are undergoing µs-ms conformational exchange throughout the mIL-2 structure. Thus, in agreement with our allosteric network model of mIL-2, stabilization of the closed/uncapped conformation through ligand binding reduces dynamic exchange at the hydrophobic core of mIL-2, as the conformational equilibrium is skewed toward the closed state. The observation of residual exchange at the CPMG time scale suggests that sampling of the open state is still permitted by the presence of the ligand, albeit with a much lower occupancy.

**Figure 5.**
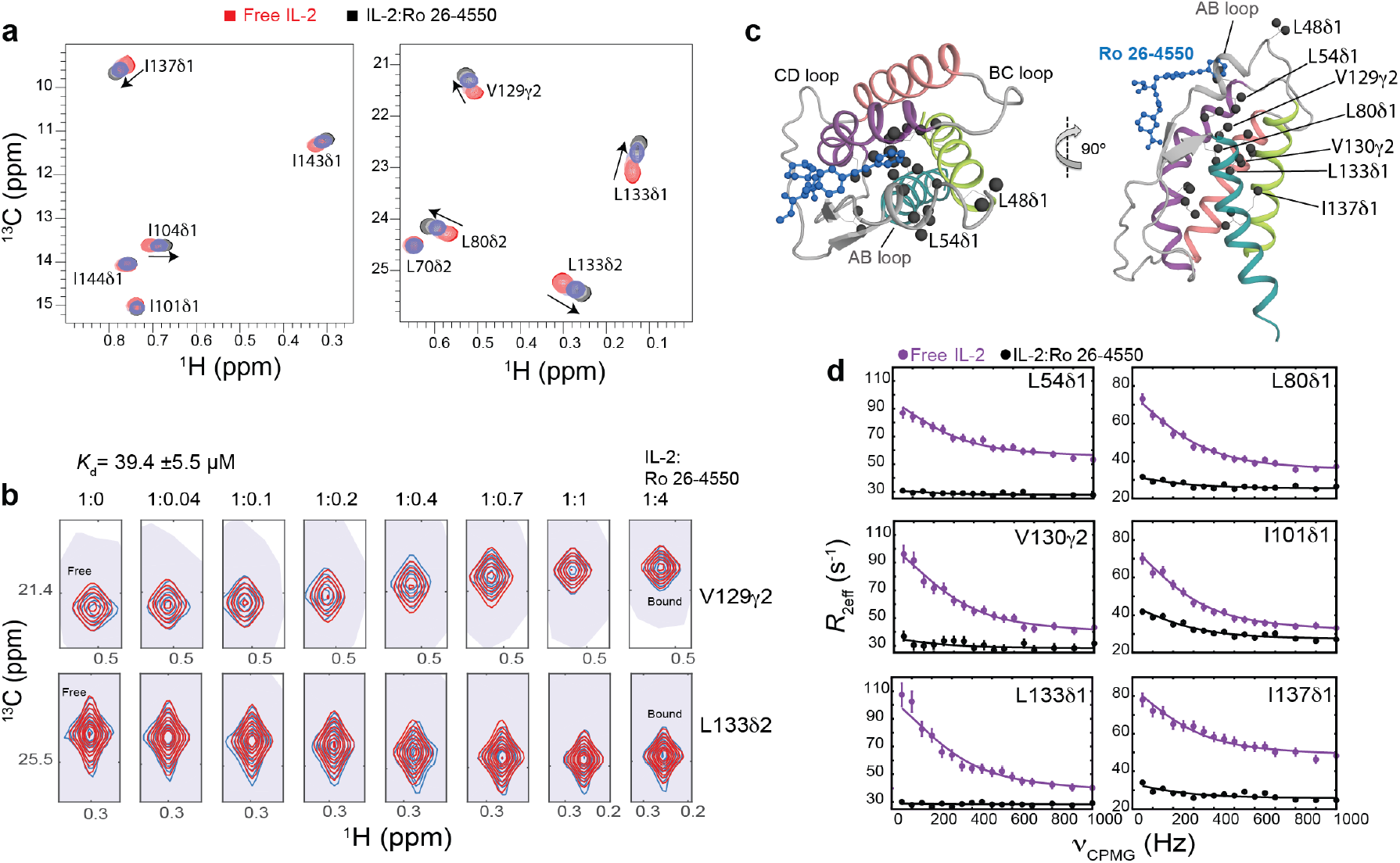
Small molecule binding at the AB loop quenches dynamics of mIL-2. (**a**) 2D ^1^H,^13^C-HMQC spectra of ILV-methyl labeled mIL-2 in the free state (red), and with increasing concentrations of Ro 26-4550 inhibitor (1:4 molar ratio, shown in black). Data were recorded at 800 MHz, 25 °C. (**b**) NMR line shape analysis for the V129γ_2_ and L133δ_2_ methyl resonances using TITAN (see methods), with indicated equilibrium dissociation constant and errors propagated from the spectral S/N. Recorded NMR spectra are shown in blue, with simulated line shapes in red. Ratios of mIL-2 to inhibitor are indicated in each panel. (**c**) Two views showing the mapping of residues undergoing significant chemical shift perturbations (CSPs) onto the ribbon representation of mIL-2. Methyl groups with marked CSPs (>0.05 ppm) are shown as black spheres. The inhibitor is shown as a blue ball-and-stick diagram on the overlaid hIL-2 complex structure (PDB: 1M48). (**d**) ^13^C single quantum CPMG relaxation dispersion profiles in the absence (purple) and presence of Ro 26-4550 at saturating concentration (black) for selected residues in the AB loop (L54δ_1_), B Helix (L80δ_1_), C Helix (I101δ_1_) and D Helix (V130γ_2_, L133δ_1_, Ι137δ_1_). CPMG experiments were performed at a ^1^H field of 600 MHz and 25 °C.

## DISCUSSION

The unique conformational plasticity of IL-2 appears to play an important role in targeting the IL-2 receptor signaling axis in both agonist and antagonist mode. However, this plasticity also presents an opportunity to target or otherwise manipulate the conformational landscape of IL-2 for drug discovery, but more detailed insights are needed for actionable value. Evidence for a functional role of such conformational plasticity was provided in previous studies focusing on both human and mouse IL-2 model system. These studies employed small molecule binding^6–7^, antibody engineering^9, 12^ or directed mutagenesis^8^ to induce minor perturbations in the cytokine’s core α-helical fold, leading to distinct immunomodulatory functions. Specifically, elucidating a link between specific allosteric effects on the IL-2 structure, induced by the JES6-1 and S4B6 antibodies, with the targeting of opposing immune cells types expressing different levels of receptor subunits *in vivo* has provided a solid foundation for immune modulation through the stabilization of different IL-2 conformational states. Conversely, the high-throughput screening of hIL-2 libraries in yeast led to the identification of a variant, termed “super-2”, comprising a helix that is locked in a conformation which is primed for interactions with the IL-2Rβ subunit, thereby leading to preferential expansion of cells lacking IL-2Rα^8^. These results were leveraged to discover a human T_reg_-based Ab, akin to JES6-1, of significant therapeutic potential^11^. Thus, given the sequence and structure conservation of the IL-2 fold across species, the results established for the mouse IL-2 paradigm directly relate to human IL-2, whose use as a multi-faceted protein therapeutic (Proleukin) has been hampered by an incomplete understanding of its precise functional properties. Therefore, a detailed characterization of the relevant conformational states and their dynamic interconversion process in solution would provide a roadmap for drugging IL-2, through a range of approaches.

Our methyl-based NMR measurements from two complementary methodologies, CPMG and CEST, provide strong support that free mIL-2 is highly dynamic at the µs-ms timescale and samples an excited state via a concerted conformational transition of the AB loop and core of the structure. Functionally relevant excited states have been previously described for a range of protein and nucleic acid systems^27^, and are typically separated by relatively low energy barriers from the ground state. The inherent plasticity of the AB loop and allosteric crosstalk with the hydrophobic core suggested an avenue for biasing the conformational equilibrium by perturbing key features present in the excited state conformation. Based on our analysis of crystallographically observed snapshots of IL-2, we hypothesized that the excited state might share similar features with the JES6-1 antibody-bound structure with respect to a stabilizing C-capping motif, and tested this hypothesis through a detailed dynamic, binding and functional characterization of a designed Ala mutant. In fact, our data demonstrate that the structural basis for such functional complexity may be surprisingly simple: the global dynamic motions of mIL-2 are induced by a conserved C-capping hydrogen-bond network, and can be switched on and off by a single point mutation (Fig. 3). Taken together, out results show that the IL-2 energy landscape is highly malleable, and establish a paradigm for controlling IL-2 cytokine activity though designed changes in protein dynamics.

Analysis of our NMR, ITC and functional data in the context of the available IL-2 co-crystal structures suggests a general scheme for molecular recognition between IL-2 and its binding partners (Fig. 6). According to this model, free mIL-2 exists in a pre-equilibrium between closed (major form, 94%) and open conformations (minor form, 6%), with the AB loop and the hydrophobic core engaged in a cooperative motion on the µs-ms timescale. In the closed/uncapped (productive) state, the AB loop is in a conformation that is primed to interact with the IL2-Rα receptor. Formation of a C-capping hydrogen network locks the AB loop in an open (autoinhibited) state, in which the loop adopts a conformation that is optimal for interactions with the JES6-1 antibody (Fig. 6 **left**). Disruption of the C-capping hydrogen bond network by mutation or small molecule binding shifts the equilibrium towards the closed state, leading to a reduction in Ab binding by 3 orders of magnitude (Fig. 6 **right**). While the “closed” and “open” conformations mediate high-affinity complex formation with the receptor and Ab, respectively, the binding process is likely achieved via the formation of an initial encounter complex, where additional induced structural adaptations follow the selection of pre-existing conformational states. This model is consistent with the majority of protein-ligand binding examples, where both processes contribute to complex formation^31^.

**Figure 6.**
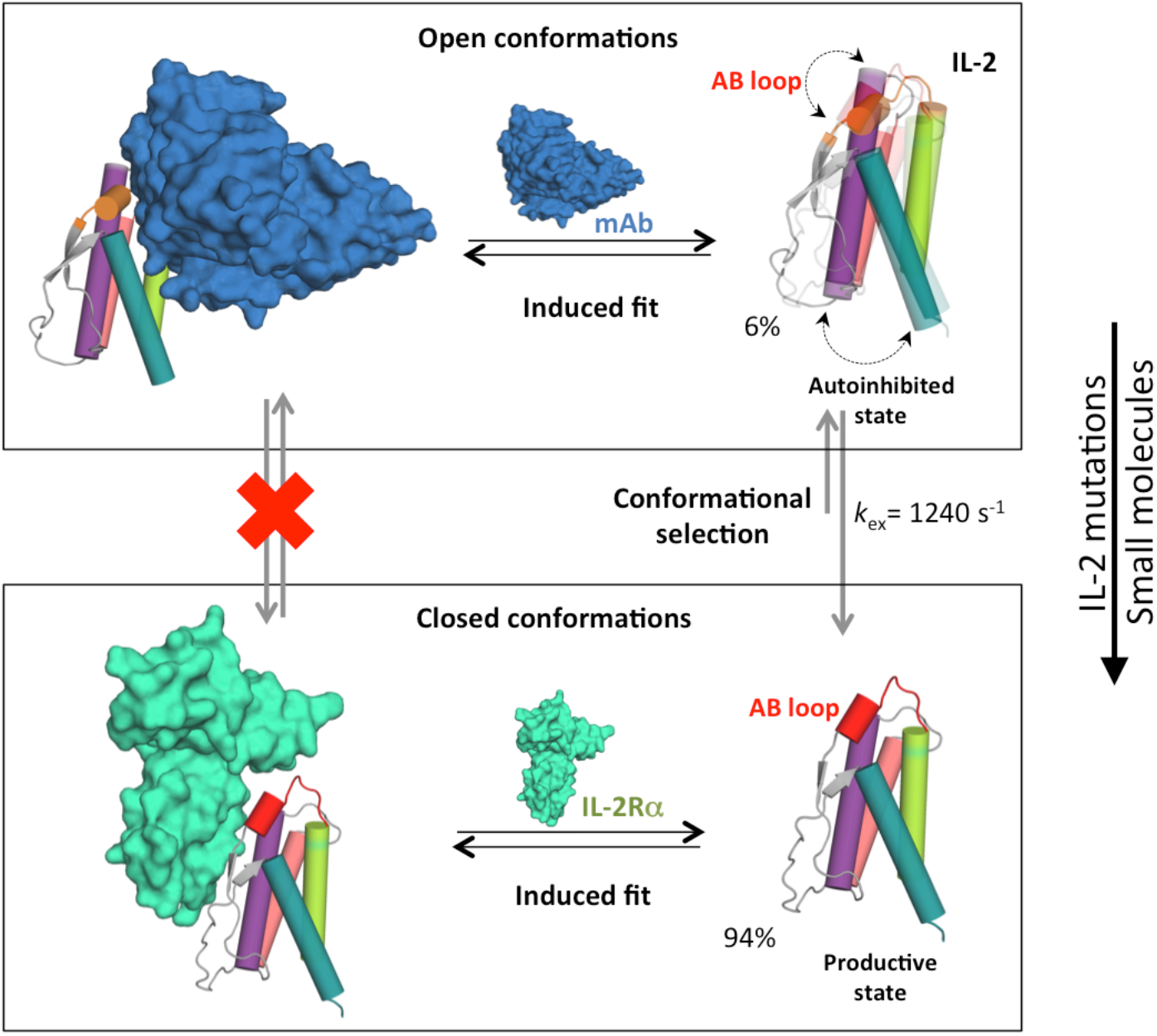
Conformational “priming” of free mIL-2 drives high-affinity complex formation. A global conformational transition enables mIL-2 to sample two distinct states that are recognized by IL2-Rα and JES6-1 via additional induced fit steps. In the closed conformation, the polar face of the AB loop including R52, is primed to interact with the IL2-Rα receptor. Formation of a C-capping hydrogen bond network by R52 locks the AB loop in an autoinhibitory, open conformation with high affinity for the JES6-1 Ab. As shown in Figures 3 and 4, point mutations or binding of small molecules can shift the equilibrium to favor the closed conformation, with measureable functional effects (right). Representative exchange parameters and excited-state populations obtained from our NMR data for mIL-2 are indicated.

A consequence of allosteric crosstalk in the IL-2 fold is that the protein becomes globally sensitive to mutations. Point mutations can induce long-range effects, which may explain the high level of sequence conservation of IL-2 residues removed from the primary receptor recognition sites. Further examples of such long-range modulation are provided by engineered IL-2 agonists. IL-2 superkine (also denoted as “super-2”) functions independently of IL-2Rα and has enhanced binding to IL-2Rβ in spite of five of the six mutations being clustered on the B-C loop and within the C helix core, removed from the IL-2Rβ primary binding site. Instead, a cluster of hydrophobic substitutions (F80, V85 and V86) induce a subtle change in C helix orientation leading to an increased affinity for IL-2Rβ. Super-2 exerts a more potent expansion of tumor-specific cytotoxic T cell populations, and a reduced expansion of T_reg_ cells^8^. Moreover, Super-2 can be used as a platform for additional mutations that inhibit binding to γc receptor, and consequently heterodimerization of IL-2Rβ and γc, thereby acting as a potent antagonist^13^. Conversely, to bypass allosteric effects present in the IL-2 fold, a *de novo* design approach was used to generate a mimetic of IL-2/IL-15 showing a new α-helical fold with a well-packed hydrophobic core, denoted Neoleukin-2/15 (Neo-2/15)^16^. This synthetic molecule retains the binding surfaces for the IL-2Rβ/γc receptors while also alleviating the requirement of IL-2Rα/IL-15Rα for formation of a productive signaling complex. Neo-2/15 exhibited enhanced anti-tumor activity in mouse models, was more stable, and lacked detectable immunogenicity^16^. These studies highlight the potential for modulating IL-2 function through different design approaches, showing varying levels of responsiveness to regulatory molecules and conditions at the vicinity of the membrane signaling complex.

In summary, our results highlight the role of conformational plasticity at conserved features of the IL-2 fold, acting locally and allosterically to form an “excited” state with discrete functional properties. Our NMR-based characterization offers a window to access otherwise hidden states of the energy landscape that can be then exploited to fine-tune the affinity and selectivity for different IL-2 binding partners towards engineering novel cytokine functions. For example, our established allosteric coupling between the receptor binding sites to the hydrophobic core of the structure presents an opportunity to alter IL-2 function through the introduction of core mutations and induction of allosteric effects without directly affecting the receptor/Ab binding surfaces. Given that the IL-2 fold presents a limited surface area with overlapping binding epitopes, our findings have important practical utility from an engineering and synthetic biology perspective.

## METHODS

### mIL-2 NMR sample preparation, backbone and methyl assignments

The sequence encoding hexahistidine-tagged mouse IL-2 (mIL-2, amino acids 1-149) was cloned into the pMAL vector with an N-terminal maltose-binding protein (MBP) followed by a 3C protease site. U-[^15^N,^13^C,^2^H]-labelled mIL-2 was overexpressed in *Escherichia coli* BL21(DE3) cells in M9 medium in ^2^H_2_O containing 2 g l^-1^ ^2^H^13^C glucose (Sigma #552151) and 1 g l^-1^ ^15^NH_4_Cl. Selective methyl labelling with Ile-δ_1_-[^13^CH_3_], Leu-δ-[^13^CH_3_/^12^C^2^H_3_], Val-γ-[^13^CH_3_/^12^C^2^H_3_], Met-ε_1_[^13^CH] referred to as MILV*, was achieved by the addition of appropriate precursors (ISOTEC Stable Isotope Products (Sigma-Aldrich) as detailed previously^34–35^. MILV-methyl (Ile ^13^Cδ1; Leu ^13^Cδ1/^13^Cδ2; Val ^13^Cγ1/^13^Cγ2; Met-^13^Cε_1_) U-[^15^N, ^2^H]-labelled mIL-2 was prepared in M9 medium in ^2^H_2_O, supplemented with 2 g l^-1^ ^2^H^12^C glucose (Sigma #552003) and 1 g l^-1^ ^15^NH Cl. Both MILV* and MILV were induced at OD_600_ ≈ 0.7 by addition of 1 mM IPTG and expression was performed for 20 h at 23 °C. Protein in the periplasmic compartment was isolated by osmotic shock and purified by nickel-nitrilotriacetic acid (Ni-NTA) (Qiagen) affinity chromatography and via size-exclusion chromatography on a Superdex-75 column (GE Healthcare) in HEPES-buffered saline (HBS, 150 mM NaCl in 20 mM HEPES pH 7.0). NMR experiments were recorded at temperature of 25 °C using 14.0 and 18.8 T magnetic field, cryoprobe-equipped Varian and Bruker spectrometers, respectively. Both MILV* and MILV IL-2 samples contained 400 to 600 µM protein in 150 mM NaCl, 20 mM HEPES pH 7.0, 0.01% NaN_3_, in 90% H_2_O/10% D_2_O. The backbone resonance assignments were obtained using a suite of TROSY-readout triple-resonance experiments^36^ (HNCO, HNCA and HNCB) and subsequently extended to the Ile, Leu and Val sidechain methyls using a 3D HMCM[CG]CBCA methyl out-and-back experiment^37^ recorded on the MILV* IL-2 sample. MILV sidechain methyl assignments were validated using methyl-to-methyl NOEs obtained from 3D H_M_-C_M_H_M_ SOFAST NOESY and 3D C_M_-C_M_H_M_ SOFAST NOESY experiments (Rossi et al., 2016). For the 3D H_M_-C_M_H_M_ SOFAST NOESY experiment, acquisition parameters were 40, 80, 1,024 complex points in the ^1^H_M_, ^13^C_M_, ^1^H_M_ dimensions with corresponding acquisition times of 25, 10 and 80 ms with 4 scans/FID. For the 3D C_M_-C_M_H_M_ SOFAST NOESY acquisition parameters were 52, 32, 1,024 complex points in the ^13^C_M_, ^13^C_M_, ^1^H_M_ dimensions with corresponding acquisition times of 13, 8 and 80 ms with 8 scans/FID. Backbone amide and sidechain methyl assignments were cross-validated using methyl-to-amide NOEs obtained from 3D H_N_-C_M_H_M_ SOFAST NOESY experiments^38^. The acquisition parameters were 64, 32 and 1,024 complex points in the ^1^H_N_-^13^C_M_, ^1^H_M_ dimensions with corresponding acquisition times of 15, 11 and 80 ms with 8 scans/FID. All 3D SOFAST NOESY experiments were recorded at 800 MHz, 25 °C on MILV-methyl-labelled mIL-2 samples using a recycle delay of 0.2 s and NOE mixing time of 300 ms. Assigned NOEs were cross-validated based on the Rosetta homology-based model of mIL-2 (using as a template PDB ID 1M47). In this manner, a set of complete assignments was obtained for the methyl groups of MILV (60) probes. All spectra were processed with NMRPipe^39^ and analyzed with CcpNMR program^40^.

### Stereospecific isotopic labeling

A specifically methyl-labeled acetolactate precursor (2-[^13^CH], 4-[^2^H] acetolactate) was obtained through deprotection and exchange of the protons of the methyl group in position 4 of ethyl 2-hydroxy-2-(^13^C)methyl-3-oxobutanoate (FB reagents) achieved in D_2_O at pH 13 (24). Typically, 300 mg of ethyl 2-hydroxy-2-(^13^C)methyl-3-oxobutanoate was added to 24 mL of a 0.1 M NaOD/D_2_O solution. After 30 min, the solution was adjusted to neutral pH with DCl and 2 mL of 1 M TRIS pH 8 in D_2_O was added. For the production of highly deuterated [U-^2^H], I-[^13^CH]δ1, L-[^13^CH]proS, V-[^13^CH]proS WT 3 3 3 mIL-2 samples, 300 mg/L of 2-[^13^CH], 4-[^2^H] acetolactate, prepared as described above, 3 3 was added 1 h prior to induction (OD_600_ ≈ 0.55). 40 min later (i.e. 20 min prior to induction), 3,3-[^2^H],4-[^13^C]-2-ketobutyrate (SIGMA #589276) was added to a final concentration of 60 mg/L. Protein was induced at OD_600_ ≈ 0.7 by addition of 1 mM IPTG and expression was performed for 20 h at 23 °C.

### SEC MALS

Absolute molecular weight calculations were obtained by static light scattering in-line with size exclusion chromatography using a Wyatt Optilab T-rEX refractometer and mini DAWN Treos multiangle light scattering system at 4 °C. mIL-2 protein samples (injection volume of 100 µL at 12mg/mL) were run at a 0.5 mL/min flow rate on a Superdex 200 10/300 GL gel filtration column (GE Healthcare) in a running buffer of 20 mM HEPES (pH 7.0), 150 mM NaCl. Protein concentrations were monitored by a refractometer and light scattering directly after the gel filtration column. Absolute molecular weights were determined using ASTRA version 6.0 (Wyatt Technologies).

### Site-directed mutagenesis

pMAL-mIL-2 derivative plasmids carrying mutations were amplified using primers containing the mutations of interest and appropriate reverse primers using a Phusion polymerase (New England Biolabs) according to the manufacturer’s recommendations. Template DNA was removed by Dpn I treatment, and transformed into *E. coli* DH5α strain. The introduced mutations and the absence of secondary mutations were verified by sequencing of plasmid DNA. Plasmids were transformed into *E. coli* BL21(DE3) strain.

### mIL-2Rα receptor and JES6-1 scFv antibody samples preparation

mIL-2Rα (amino acids 1-213) ectodomain was secreted and purified using a baculovirus expression system, as previously described^41^. Recombinant JES6-1 single-chain Fv (scFv) was expressed and purified in a baculovirus expression system as described earlier^15^. All proteins were purified to >98% homogeneity with a Superdex 200 sizing column (GE Healthcare) equilibrated in 150 mM NaCl, 10 mM HEPES pH 7.3. Purity was verified by SDS-PAGE analysis.

### STAT5 signaling

CTLL-2 cells (AATC TIB-214), a murine IL-2 dependent T cell line, were maintained in complete RPMI (RPMI 1640-glutaMAX supplemented with 10 % fetal bovine serum, non-essential amino acids, sodium pyruvate, 15 mM HEPES and penicillin-streptomycin supplemented with 1000 IU/mL recombinant mouse at 37 °C with 5 % CO_2_. Cells were rested in RPMI for 16 hours prior to signaling analysis. For STAT5 signaling, 2-3 x10^5^ cells were cultured in 100 µL RPMI with cytokine or cytokine:antibody mixture (2:1) in a 96 well plate prior to fixation with 1.6 % paraformaldehyde for 10 min at room temperature. Cells were permeabilized with 100 % ice-cold methanol and stored at −20 °C prior to staining. Cells were washed twice with FACS buffer (PBS pH 7.2, 2 % FBS, 2 mM EDTA) and stained with 1:100 Alexa Fluor 647 conjugated anti-STAT5 pY694 (BD) for 1 hour at room temperature. Mean fluorescence intensity (MFI) was monitored using a CytoFLEX flow cytometer (Beckman Coulter).

### mIL-2Rα and JES6-1 NMR chemical shift mapping

2D ^1^H-^13^C SOFAST-HMQC were acquired on 100-200 µM Iδ_1_-^13^CH_3_, L, V proS methyl-labelled mIL-2 in the free state and in the bound state in a 1:1 molar complex with IL-2Rα or JES6-1 at 800 MHz, 25 °C. Acquisition parameters were 256 and 1,024 complex points in the ^13^C_M_, ^1^H_M_ dimensions with corresponding acquisition times of 58 ms and 80 ms using a relaxation delay of 0.2 s with 8 scans/FID in the free state and 32 scans/FID in the bound state. The change in chemical shift (in p.p.m.) between the free and IL-2Rα or JES6-1 bound state of mIL-2 I(LV)proS-methyls was determined using the equation Δδ^CH3^ = [1/2 (Δδ^2^H + Δδ^2^C/4)]^1/2^. To confirm the assignments of mIL-2 I(LV)proS-methyl peaks that shifted upon IL-2Rα binding, an additional 3D C_M_-C_M_H_M_ SOFAST NOESY was acquired on labelled mIL-2 in a 200 µM 1:1 complex with IL-2Rα using 45, 30, 1,024 complex points in the ^13^C_M_, ^13^C_M_, ^1^H_M_ dimensions with corresponding acquisition times of 11, 7.5 and 80 ms and 48 scans/FID. The assignments of mIL-2 I(LV)proS-methyl peaks that shifted upon JES6-1 binding were confirmed by a 3D C_M_-C_M_H_M_ SOFAST NOESY using 45, 20, 1,024 complex points in the C_M_-C_M_H_M_ dimensions with corresponding acquisition times of 10, 4.5 and 80 ms and 32 scans/FID.

### Small molecule NMR titration

The titration of the small molecule Ro 26-4550 (Tocris Bioscience) onto ILV methyl-labeled WT mIL-2 was performed on a 350 µM sample in NMR buffer (150 mM NaCl, 20 mM HEPES pH 7.2, 2.5% deuterated DMSO) at the following mIL2-Inhibitor ratios: 1:0, 1:0.04, 1:0.1, 1:0.2, 1:0.4, 1:0.7, 1:1, 1:1.2 and 1:4, with 2D ^1^H-^13^C SOFAST HMQC spectra as a readout. The mIL-2 sample was saturated (∼97%) by the addition of a 4-fold excess of inhibitor. A small dilution of the protein sample occurred over the course of the titrations due to addition of the inhibitor solution aliquots. The compound was prepared as a 1mM stock solution in NMR buffer containing 2.5% deuterated DMSO. The change in chemical shift (in p.p.m.) between the free and Ro 26-4550 bound state of mIL-2 ILV-methyls was determined using the equation as described above. Data were processed with 4 Hz and 10 Hz Lorentzian line broadening in the direct and indirect dimensions, respectively and fit using a two-state model in TITAN^42^ with bootstrap error analysis of 100 replicas. A total of eight NMR peaks were used for the global fitting procedure.

### Methyl CPMG relaxation dispersion experiments

Methyl single-quantum ^13^C CPMG relaxation dispersion experiments (Lundstrom et al., 2007) were recorded on highly deuterated ILV-methyl labeled WT and R52A mIL-2 samples (both at 400 µM protein concentration) at field strengths of 14.0 T and 18.8 T, at 25 °C, using Varian and Bruker spectrometers, both equipped with a cryogenically cooled probe. The CPMG data set was acquired as pseudo 3D experiments with a constant relaxation time period T_relax_ of 20 ms and with 18 CPMG pulse frequencies ν_CPMG_ = 1/(2τ) ranging from 50 to 1000 Hz, where τ is the delay between the consecutive 180° refocusing pulses in ^13^C CPMG pulse-train. Relaxation dispersion profiles R_2,eff_(ν_CPMG_) were calculated from peak intensities (I) recorded at different CPMG frequencies ν_CPMG_ using the following equation: R_2,eff_(ν_CPMG_) = −1/T_relax_ln(I/I_0_), where I is signal intensity in the spectra collected at T_relax_ = 20 ms, I_0_ is signal intensity in the reference spectrum recorded at T_relax_ = 0. An interscan delay of 1.5 s was used with 24, 32 or 36 scans/FID, giving rise to net acquisition times between 40-58 h for a complete pseudo-3D data set. All data were processed using NMRpipe^39^ and peak intensities were picked using CCPN^40^. The error was determined from the noise level of the spectra. The variation in R_2,eff_ with ν_CPMG_ was fit to a two-state model of chemical exchange based on the Bloch-McConnell equations, to extract values of exchange parameters (*p*_B_, *k*_ex_=*k*_AB_+*k*_BA_), as well as ^13^C chemical shift differences for nuclei interconverting between pairs of states. The software CATIA^43^ was used to fit the data. Initially, global fits included 6 profiles for WT mIL-2 at two magnetic fields (L54δ1, L80δ2, V83γ1, L84δ1, I101δ1, L133δ1). The fitting was performed by minimizing the function χ^2^ as previously described^44^. The group fit of selected residues was performed if the χ^2^_Group_/χ^2^_Local_ was less than 2.0. As a second step, 14 profiles (only from 14.0 T magnetic field) were used to obtain |Δω| values calculated from per-residue and global fits with *k*_ex_ and *p*_B_ fixed to the values obtained in global data fits. For R52A mIL-2, global fits included 8 profiles at two magnetic fields (L60δ2, L80δ2, L84δ1, V109γ1, V129γ1, V130γ2, L133δ1, I137δ1).

### ^13^C CEST

^13^C CEST experiments^45^ were recorded on highly deuterated ILV-methyl labeled WT and R52A mIL-2 samples (both at 400 µM protein concentration) at field strength of 18.8 T at 10°C with an exchange period (T_CEST_) of 300 ms and using a pair of *B*_1_ fields, 16.2 and 32.4 Hz. For WT mIL-2, 135 data sets were obtained for *B*_1_ = 16.2 Hz, with offsets ranging from 16 to 26 ppm (82 spectra, 25 Hz steps) and from 5.3 to 12.8 ppm (53 spectra, 30 Hz steps); 52 spectra were recorded from 16 to 26 ppm in 40 Hz increments for *B*_1_ = 32.4 Hz. For R52A mIL-2, 135 data sets were obtained for *B*_1_ = 16.2 Hz, with offsets ranging from 16 to 26 ppm (82 spectra, 25 Hz steps) and from 5.3 to 12.8 ppm (53 spectra, 30 Hz steps). Each 2D data set comprised of 60 (^13^C) x 512 (^1^H) complex points (28 ms, 64 ms), 16 scans/FID and a repetition delay of 2.0 s corresponding to a total measuring time of 88 (*B*_1_ =16.2 Hz) and 63 (*B*_1_ =32.4 Hz) h for each 2D series. Experiments were processed using NMRpipe^39^ and CEST profiles were generated as the ratio in intensities of peaks in spectra acquired with and without the T_CEST_ period vs the position of the low power *B*_1_ field. Uncertainties in I/I_0_ were determined from the scatter in the baseline of CEST profiles where no intensity dips are present (typically, n > 30). Weak *B*_1_ fields applied during T_relax_ were calibrated according to the procedure^46^. The CEST profiles were analyzed using the program Chemex (https://github.com/gbouvignies/chemex), which numerically propagates the Bloch– McConnell equation as described^47^. Initially, data sets were analyzed simultaneously and all residues were included in the analysis, including those for which well-resolved excited state dips were not obtained. For the two-site exchange model, only residues that showed distinct major and minor dips or asymmetry were fit globally (nine residues) including the initial *k*_ex_ and *p*_b_, and residue-specific Δω values. As a third step, we fixed the global exchange parameters (*k*_ex_ and *p*_b_) and re-fitted all residues.

### Sidechain rotamer space analysis in mIL-2

We analyzed sidechain rotamer space of buried residues of mIL-2 using a customized software tool (*manuscript in preparation*) (**Extended Data Fig. 7a**). This software takes as input, a protein structure and then explores the sidechain rotamer space of the residues of interest. Before examining the space of rotamers, sidechains of the input structure are removed. Every sidechain rotamer of a residue obtained from a backbone dependent rotamer library is plugged in to check for a steric clash with its or its neighbors’ backbones. All the rotamers that result in clashes with backbone are eliminated. Next, rotamers of neighboring residues are considered in pairs; clashing rotamer pairs are retained as constraints for subsequent step. The information about independent and pairwise rotamers is utilized towards an exhaustive global check phase that eliminates all the rotamers that are incompatible in all the valid combination of rotamers in a structure (**Extended Data Fig. 7b-d**). Finally, the reduced rotamer set for each residue is constructed and output by the tool.

### Isothermal titration calorimetry

Titrations of WT and R52A mIL-2 (40 or 96 µM) into JES6-1 (4 or 9.6 μM) were performed at 20 °C, using a MicroCal VP-ITC instrument. To minimize enthalpy of solvation effects, all experiments were performed in phosphate buffer. All protein samples were extensively dialyzed against ITC buffer (20 mM sodium phosphate, pH 7.2, 150 mM NaCl). All solutions were filtered using membrane filters (pore size, 0.22 μm) and thoroughly degassed for 20 min before the titrations. Typically, two injections of 2 μL were followed by 26 injections of 10 μL until a molar ratio of 2.0-3.0 was obtained. The dilution heats are typically small and were subtracted from the calorimetric data. Integration of the thermogram and subtraction of the blanks yielded a binding isotherm that was fitted to a one-site binding model using the MicroCal Origin 7.0 software (OriginLab Corporation) to determine the stoichiometric ratios, the dissociation constants and the changes in enthalpy.

## Acknowledgment

This research was supported by NIAID (AI2573-01) and NIGMS (1R35GM125034-01) grants to N.G.S., and by a High End Instrumentation (HIE) Grant (S10OD018455), which funded the 800 MHz NMR spectrometer at UCSC. The authors would like to acknowledge Drs Hsiau-Wei Lee and Andrew McShan for assistance with recording NMR data, and Sarvind Tripathi and Jeff Swan for assistance with SEC-MALS analysis. We thank Dr Enrico Rennella (Lewis Kay’s Group at the University of Toronto) for help with the implementation of ^13^C-Methyl CEST experiment.

## Author Contributions

V.S.P., K.C.G. and N.G.S. designed the study. K.M.J. and C.R.G. prepared IL-2Rα and JES6-1 samples and performed experiments in vitro. V.S.P. generated constructs, prepared and purified isotopically labeled proteins and performed isothermal titration calorimetry experiments. V.S.P. and N.G.S. acquired and analyzed NMR data. S.N. and N.G.S. performed sidechain rotamer space analysis. V.S.P., K.M.J, K.C.G. and N.G.S wrote the manuscript.

## Data availability

Nuclear Magnetic Resonance assignments for the free WT mIL-2, mIL-2 in complex with JES6-1 scFv antibody, mIL-2 in complex with IL-2Rα and R52A mIL-2 mutant have been deposited in the BMRB under accession numbers 27969, 27970, 27971 and 27974, respectively.

---

**Supplementary Fig. 1.**
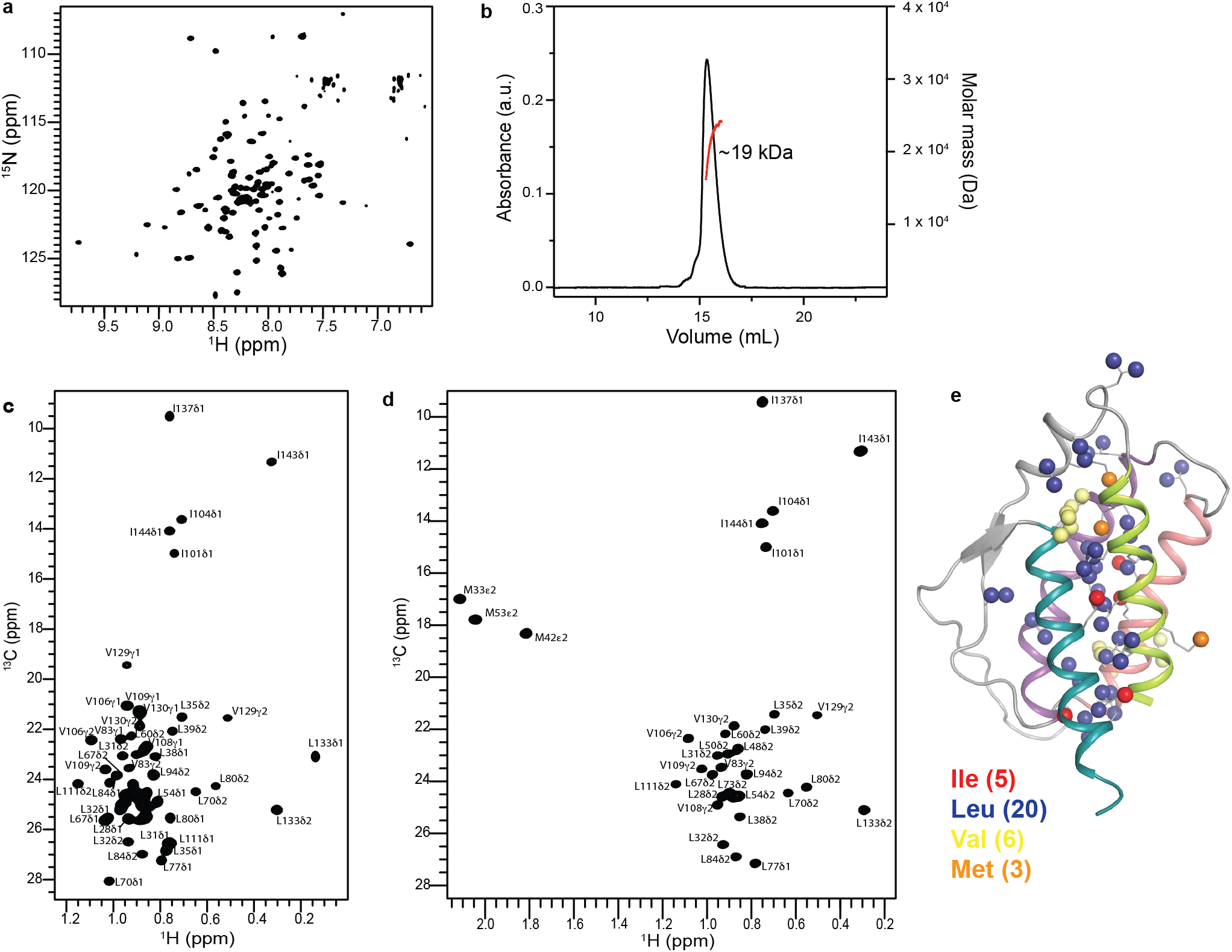
NMR spectra of free mIL-2. **(a)** ^1^H-^15^N TROSY HSQC, **(c)** ^1^H-^13^C methyl HMQC spectra of [U-^2^H,^15^N, Ileδ_1_-^13^CH_3_; Leu, Val-^13^CH_3_/^12^CD_3_] and **(d)** ^1^H-^13^C methyl HMQC spectra of [U-^2^H,^15^N, Iδ_1_-^13^CH_3_, L, V proS]-labeled mIL-2 acquired at 800 MHz, 25 °C. **(b)** MALS of free WT mIL-2 shows that the protein is monomeric in solution. **(e)** Distribution of assigned MILV methyl probes in the free mIL-2 structure. The backbone of the mIL-2 is show as in Figure 1. Color-coding: Ile, red; Leu, blue; Val, yellow; Met, orange. The numbers in parenthesis indicate the number of the corresponding residues in mIL-2.

**Supplementary Fig. 2.**
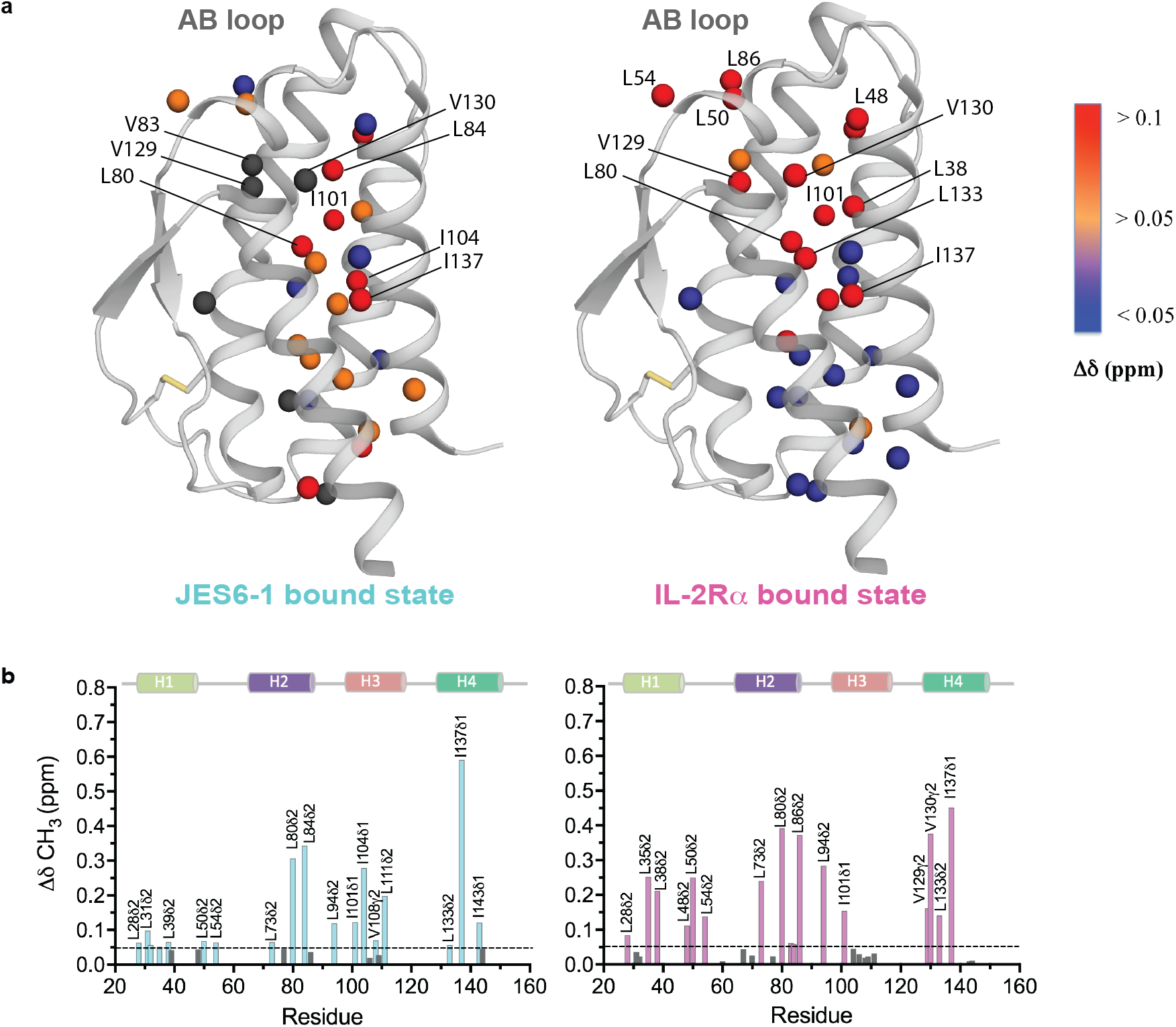
Long-range effects in mIL-2 core dynamics induced by binding to the mIL-2Rα receptor or JES6-1 immunomodulatory scFv. **(a)** Effect of IL-2Rα receptor and JES6-1 scFv on the mIL-2 structure as assessed by chemical shift perturbation. The black spheres represent missing residues of mIL-2 due to significant exchange broadening upon JES6-1 binding. Chemical shift difference (Δδ, ppm) values are mapped by continuous-scale color onto the mIL-2 structure. **(b)** Histogram of chemical shift perturbations (Δδ) as a function of IL-2 residue number. Mapping of methyl chemical shift changes on the mIL-2 structure are shown in Figure 1. CSPs were calculated as described in Methods.

**Supplementary Fig. 3.**
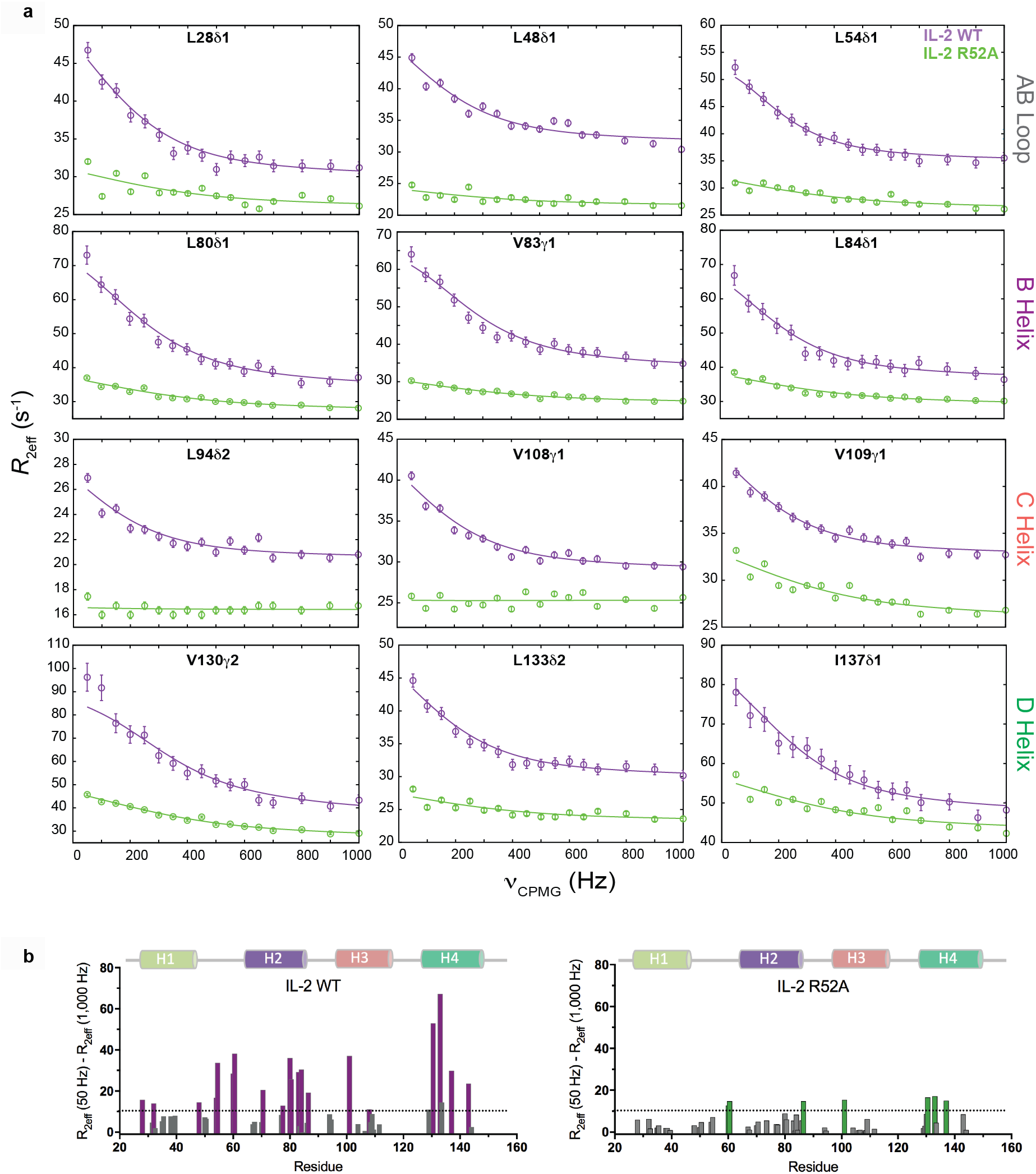
Relaxation dispersion experiments indicate that R52A mutant abrogates conformational exchange in IL-2. (**a**) Experimental relaxation dispersion profiles (circles) for residues exhibiting µs-ms timescale dynamics as measured by ^13^C SQ CPMG relaxation dispersion experiments for the WT (purple) and R52A (green) mIL-2, acquired at 600 MHz (25 °C). Solid lines represent the best fit to a global two-site exchange model. Twenty residues of WT and R52A IL-2 were used for the global analysis using CATIA program. (**b**) Plots of the *R*_ex_ contributions of the methyl groups for WT and R52A mutant. *R*_ex_ contributions were calculated from the differences between R_2eff_ (50 Hz) and R_2eff_ (1000 Hz). The methyl groups with significant *R*_ex_ contributions (> 10 Hz) are colored purple for WT and green for R52A mIL-2.

**Supplementary Fig. 4.**
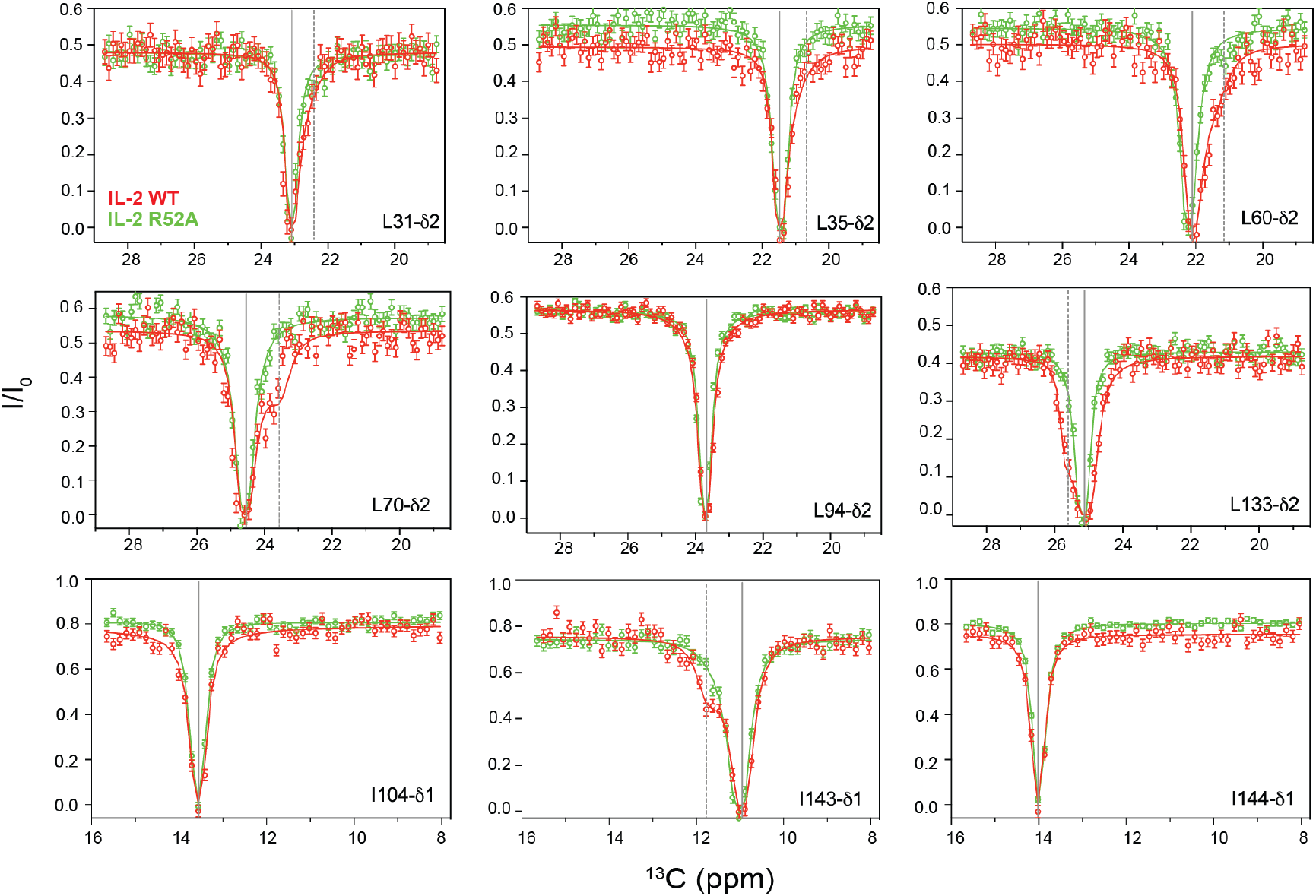
R52A mutation disrupts the conformational dynamics of mIL-2 as measured by NMR. Representative of ^13^C-CEST profiles obtained for WT (red) and R52A (green) mIL-2, recorded at a B_1_ saturation field of 16.4 Hz (T_CEST_ = 300 ms), acquired at 800 MHz and 10 °C. The chemical shift for the major state (solid lines) and minor state (doted lines) are indicated. Uncertainties in I/I_0_ for CEST data are determined from the rmsd in the baseline of the profile where no intensity dips are present (typically, n > 30). L94, I104 and I144 are examples of residues in which a minor dip were not observed.

**Supplementary Fig. 5.**
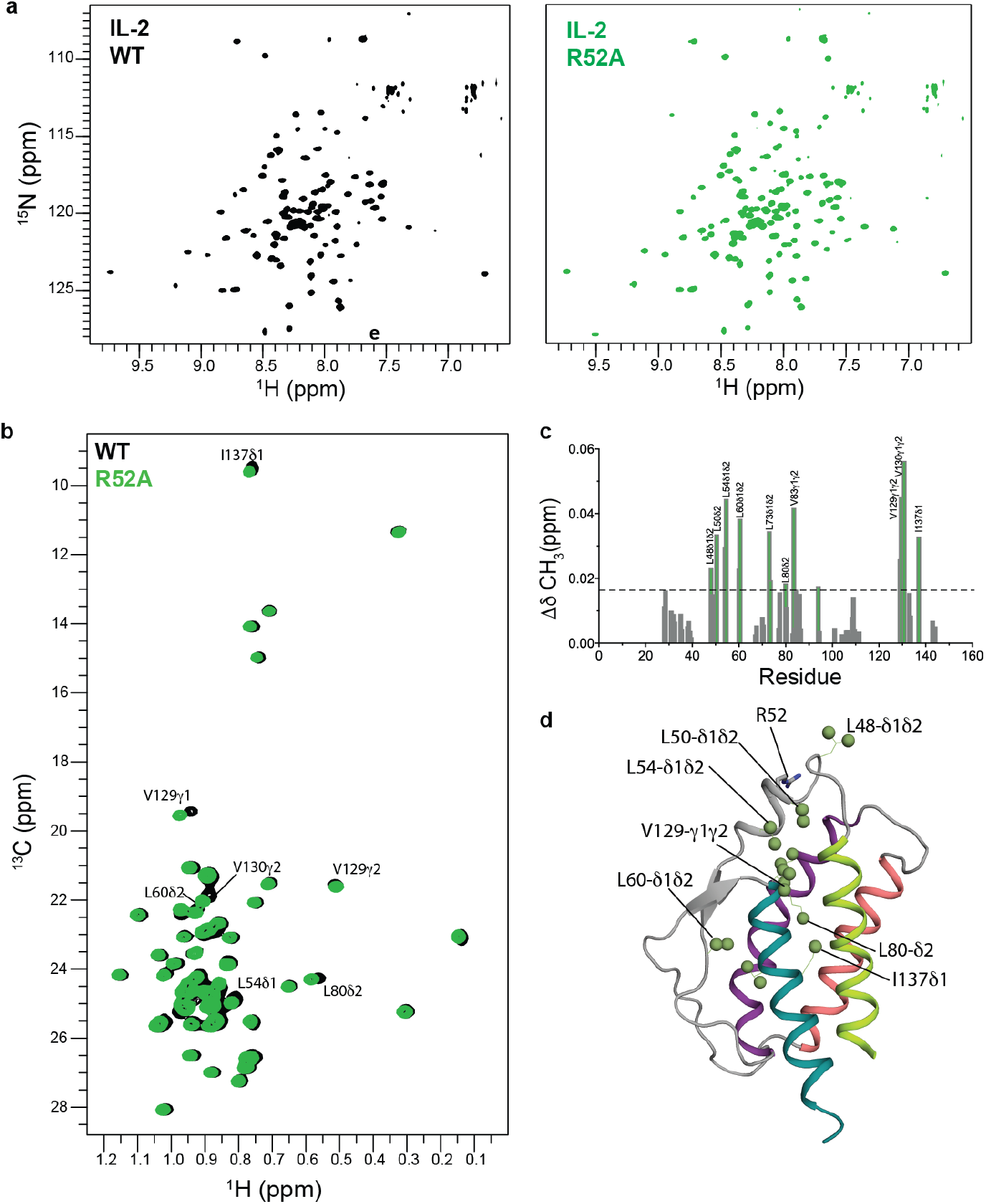
^15^N and ^13^C NMR characterization of R52A mIL-2. **(a)** Comparison of ^1^H-^15^N TROSY HSQC and (**b**) Superimposed ^1^H-^13^C-HMQC spectra of ILV-methyl labeled WT (black) and R52A (green) mIL-2, respectively. Assignments are indicated for residues with the largest chemical shift changes. Data acquired at 800 MHz, 25 °C. (**c**) Histograms of chemical shift perturbations in ILV-methyl labeled IL-2. Residues with CSPs values 1s above the average are indicated (black doted line). CSPs were calculated as described in Materials and Methods. (**d**) Mapping of the methyl groups with marked chemical shift differences onto the mIL-2 structure. The results of the effect of this mutation on the dynamics of mIL-2 and binding to JES6-1 are shown in Figure 3.

**Supplementary Fig. 6.**
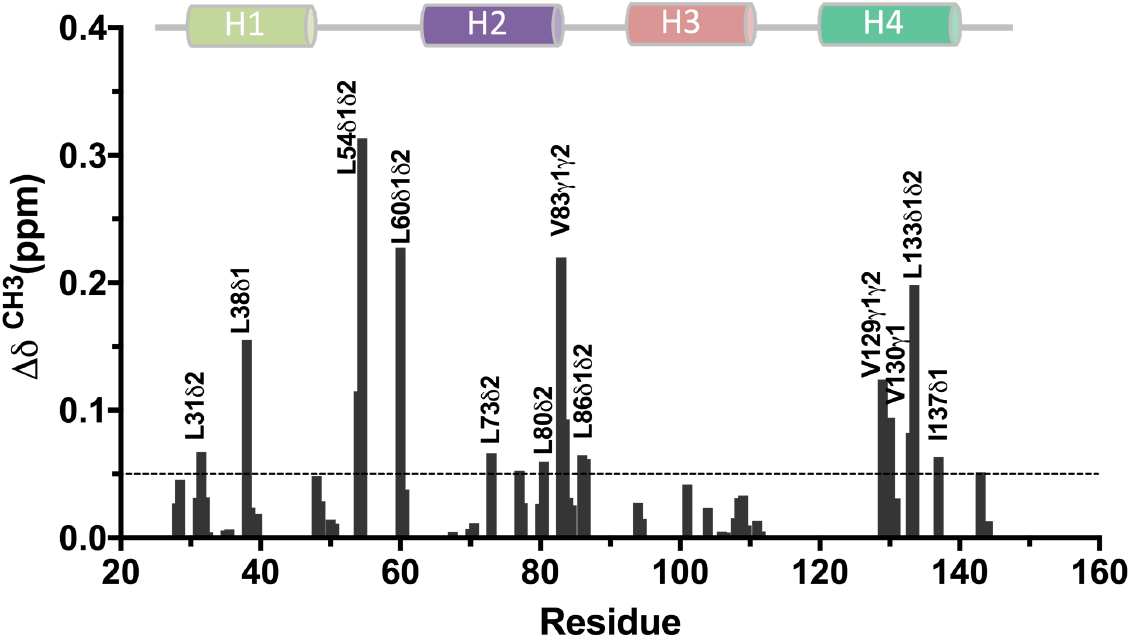
Small molecule binding induces long-range effects and quenches dynamics in mIL-2. Histogram of chemical shift perturbations (Δδ) as a function of IL-2 residue number. Mapping of methyl chemical shift changes on the mIL-2 structure are shown in Figure 5. CSPs were calculated as described in Materials and Methods

**Supplementary Fig. 7.**
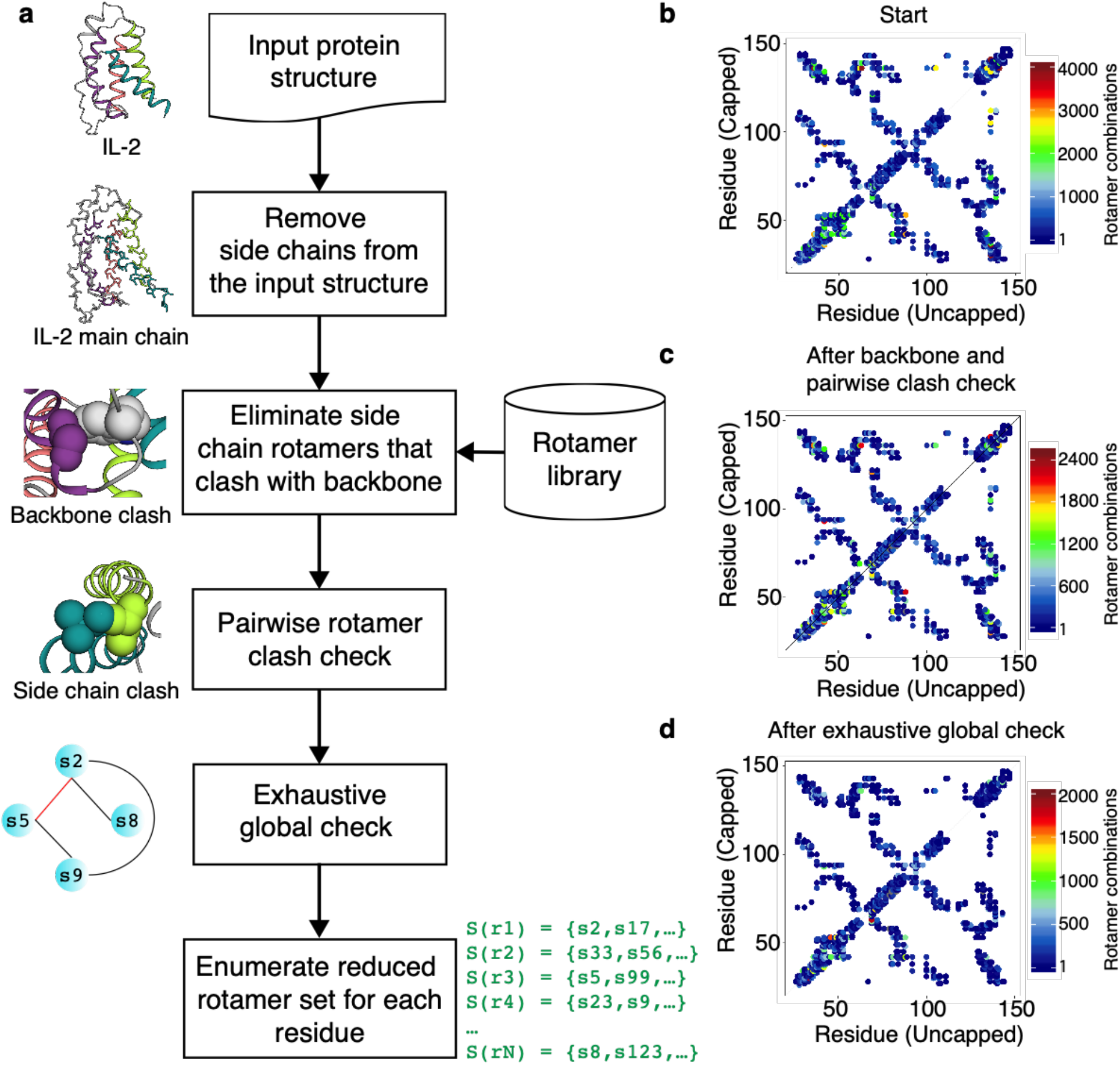
**(a)** Workflow of the method used to perform side chain rotamer space analysis. **(b)** Number of rotamer combinations between neighboring residues along the sequence of IL-2. An upper bound for number of rotamer combinations after **(c)** backbone and pairwise clash check, and **(d)** exhaustive global check between neighboring residues. Here, the upper triangular matrix shows rotamer combinations for a capped structure, whereas the lower triangular matrix for an uncapped structure.

**Supplementary Fig. 8.**
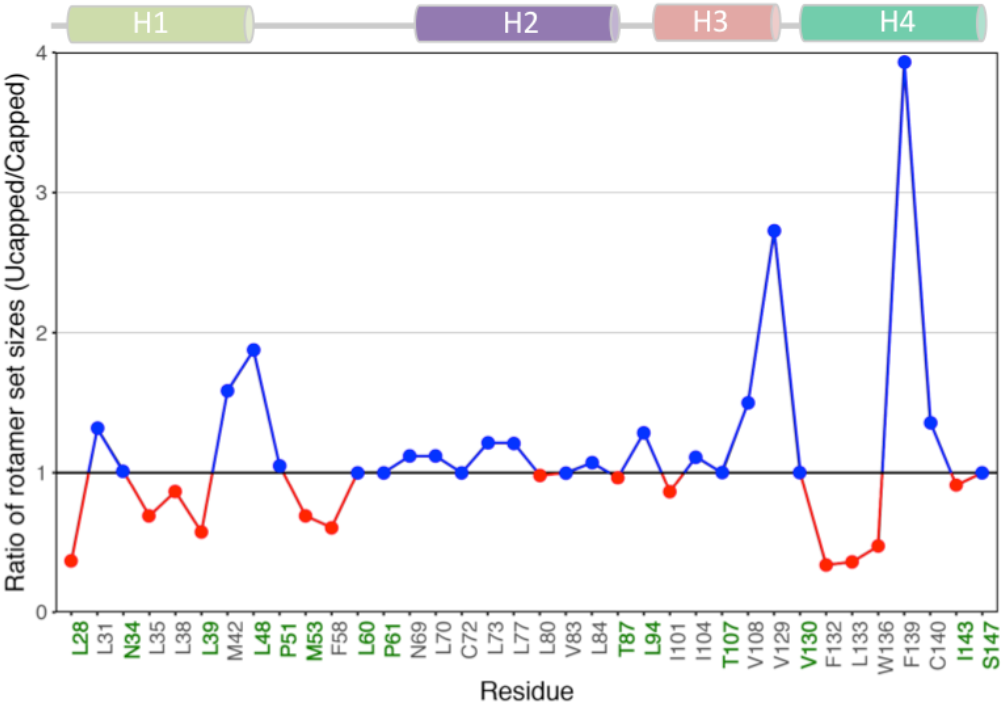
Ratio of rotamer set sizes between the closed and open mIL-2 states. Blue/red points indicate residues showing expansion/contraction of allowed rotameric states in the uncapped structure. Black are residues that are buried in both capped and uncapped structures, and green residues that are buried in one of the two structures.

**Supplementary Table 1.** Global fits of the ^13^C-CPMG relaxation dispersion curves measured at 600 and 800 MHz at 25 °C for WT mIL-2.

| Residue | $R_{\text{ex}}$<br>(600 MHz, $\text{s}^{-1}$) | $\Delta\omega$ (ppm) | $p_{\text{B}}$ (%) | $k_{\text{ex}}$ ($\text{s}^{-1}$) |
| --- | --- | --- | --- | --- |
| L28-$\text{C}\delta_1$ | 15.8 | 0.75 (0.02) | $5.8 \pm 0.4$ | $1,241 \pm 72$ |
| L35-$\text{C}\delta_2$ | 7.6 | 0.44 (0.04) | | |
| L48-$\text{C}\delta_1$ | 14.5 | 0.67 (0.01) | | |
| L54-$\text{C}\delta_1$ | 16.8 | 0.71 (0.03) | | |
| L60-$\text{C}\delta_1$ | 28.5 | 1.27 (0.01) | | |
| L60-$\text{C}\delta_2$ | 38.1 | 1.57 (0.01) | | |
| L70-$\text{C}\delta_2$ | 20.6 | 0.83 (0.04) | | |
| L80-$\text{C}\delta_1$ | 36.1 | 1.31 (0.05) | | |
| L80-$\text{C}\delta_2$ | 25.7 | 0.98 (0.04) | | |
| V83-$\text{C}\gamma_1$ | 29.3 | 1.1 (0.06) | | |
| L84-$\text{C}\delta_1$ | 30.4 | 1.02 (0.06) | | |
| L86-$\text{C}\delta_2$ | 19.2 | 0.91 (0.02) | | |
| L94-$\text{C}\delta_2$ | 6.2 | 0.42 (0.02) | | |
| I101-$\text{C}\delta_1$ | 37.1 | 1.43 (0.05) | | |
| V108-$\text{C}\gamma_1$ | 11.1 | 0.59 (0.01) | | |
| V109-$\text{C}\gamma_1$ | 8.7 | 0.54 (0.01) | | |
| V130-$\text{C}\gamma_2$ | 52.9 | 1.89 (0.01) | | |
| L133-$\text{C}\delta_1$ | 67.3 | 1.94 (0.01) | | |
| L133-$\text{C}\delta_2$ | 14.4 | 0.69 (0.02) | | |
| I137-$\text{C}\delta_1$ | 29.9 | 1.22 (0.06) | | |

**\*In order to justify group fitting, $\chi^2_{\text{group}}/\chi^2_{\text{local}} \leq 2$ where $\chi^2_{\text{local}}$ is the $\chi^2$ when a residue is locally fit and $\chi^2_{\text{group}}$ is the $\chi^2$ of the same residue when group fit.**

**Supplementary Table 2.** NOE violation analysis for the mIL-2 open and closed structures.

|  | Closed<sup>b</sup> | Open<sup>c</sup> |
| --- | --- | --- |
| **NOE distance constraint statistics** |  |  |
| Total NOE distance constraints<sup>a</sup> | 26 | 26 |
| Distance violations |  |  |
| less than 1 Å | 7 | 4 |
| between 1 and 2 Å | 1 | 2 |
| between 2 and 5 Å | 0 | 3 |
| greater than 5 Å | 0 | 7 |

<sup>a</sup> We used a strict upper distance bound up to 6 Å for the NOE restraints assigned manually.
<sup>b</sup> mIL-2 Rosetta homology model
<sup>c</sup> PDB: 4YQX

**Supplementary Table 3.**
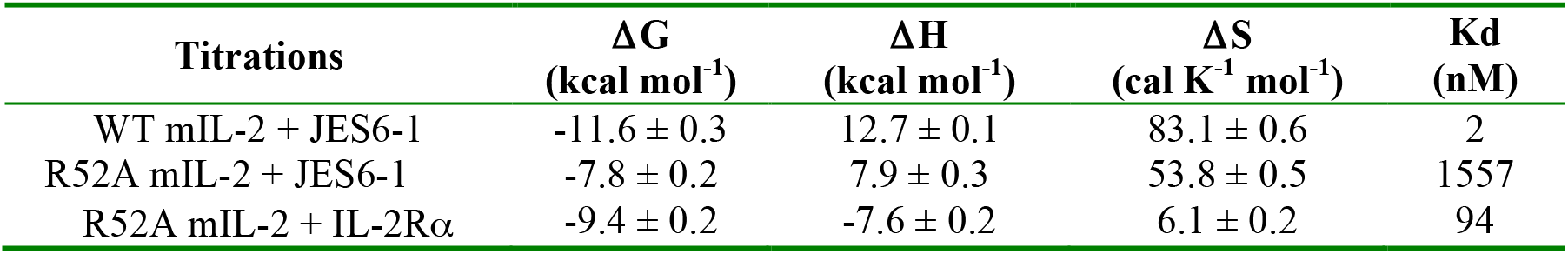
Parameters obtained from the ITC experiments measured at 20 °C.

| Titration | $\Delta G$<br>(kcal mol<sup>-1</sup>) | $\Delta H$<br>(kcal mol<sup>-1</sup>) | $\Delta S$<br>(cal K<sup>-1</sup> mol<sup>-1</sup>) | K<sub>d</sub><br>(nM) |
| --- | --- | --- | --- | --- |
| WT mIL-2 + JES6-1 | -11.6 ± 0.3 | 12.7 ± 0.1 | 83.1 ± 0.6 | 2 |
| R52A mIL-2 + JES6-1 | -7.8 ± 0.2 | 7.9 ± 0.3 | 53.8 ± 0.5 | 1557 |
| R52A mIL-2 + IL-2Rα | -9.4 ± 0.2 | -7.6 ± 0.2 | 6.1 ± 0.2 | 94 |

## REFERENCES

1. Boyman, O., Sprent, J. The role of interleukin-2 during homeostasis and activation of the immune system. Nat. Rev. Immunol. 12, 180–190 (2012).

2. Liao, W., Lin, J.-X., Leonard, W.J. Interleukin-2 at the crossroads of effector responses, tolerance, and immunotherapy. Immunity 38, 13–25 (2013).

3. Malek, T.R., Castro, I. Interleukin-2 receptor signaling, at the interface between tolerance and immunity. Immunity 33, 153–165 (2010).

4. Taniguchi, T. The IL-2/IL-2 receptor system: A current overview. Cell 73, 5–8 (1993).

5. Malek, T.R., Bayer, A.L. Tolerance, not immunity, crucially depends on IL-2. Nat. Rev. Immunol. 4, 665–74 (2004).

6. Thanos, C.D., DeLano, W.L., Wells, J.A. Hot-spot mimicry of a cytokine receptor by a small molecule. Proc Natl Acad Sci U S A. 103, 15422–7 (2006).

7. Arkin, M.R., Randal, M., DeLano, W.L., Hyde, J., Luong, T.N., Oslob, J.D., Raphael, D.R., Taylor, L., Wang, J., McDowell, R.S., Wells, J.A., Braisted, A.C. Binding of small molecules to an adaptive protein-protein interface. Proc Natl Acad Sci U S A. 100, 1603–8 (2003).

8. Levin, A.M., Bates, D.L., Ring, A.M., Krieg, C., Lin, J.T., Su, L., Moraga, I., Raeber, M.E., Bowman, G.R., Novick, P., Pande, V.S., Fathman, C.G., Boyman, O., Garcia, K.C. Exploiting a natural conformational switch to engineer an interleukin-2 ‘superkine’. Nature 484, 529–533 (2012).

9. Létourneau, S., van Leeuwen, E.M., Krieg, C., Martin, C., Pantaleo, G., Sprent, J., Surh, C.D., Boyman, O. IL-2/anti-IL-2 antibody complexes show strong biological activity by avoiding interaction with IL-2 receptor alpha subunit CD25. Proc. Natl Acad. Sci. USA 107, 2171–2176 (2010).

10. Spangler, J.B., Tomala, J., Luca, V.C., Jude, K.M., Dong, S., Ring, A.M., Votavova, P., Pepper, M., Kovar, M., Garcia, K.C. Antibodies to Interleukin-2 Elicit Selective T Cell Subset Potentiation through Distinct Conformational Mechanisms. Immunity. 42, 815–825 (2015).

11. Trotta, E., Bessette, P.H., Silveria, S.L., Ely, L.K., Jude, K.M., Le, D.T., Holst, C.R., Coyle, A., Potempa, M., Lanier, L.L., Garcia, K.C., Crellin, N.K., Rondon, I.J., Bluestone, J.A. A human anti-IL-2 antibody that potentiates regulatory T cells by a structure-based mechanism. Nat Med. 24, 1005–1014 (2018).

12. Boyman, O., Kovar, M., Rubinstein, M.P., Surh, C.D., Sprent, J. Selective stimulation of T cell subsets with antibody–cytokine immune complexes. Science 311, 1924–1927 (2006).

13. Mitra, S., Ring, A.M., Amarnath, S., Spangler, J.B., Li, P., Ju, W., Fischer, S., Oh, J., Spolski, R., Weiskopf, K., Kohrt, H., Foley, J.E., Rajagopalan, S., Long, E.O., Fowler, D.H., Waldmann, T.A., Garcia, K.C., Leonard, W.J. Interleukin-2 activity can be fine tuned with engineered receptor signaling clamps. Immunity 42, 826–838 (2015).

14. Arenas-Ramirez, N., Zou, C., Popp, S., Zingg, D., Brannetti, B., Wirth, E., Calzascia, T., Kovarik, J., Sommer, L., Zenke, G., Woytschak, J., Regnier, C.H., Katopodis, A., Boyman, O. Improved cancer immunotherapy by a CD25-mimobody conferring selectivity to human interleukin-2. Sci. Transl. Med. 8, 367ra166 (2016).

15. Spangler, J.B., Trotta, E., Tomala, J., Peck, A., Young, T.A., Savvides, C.S., Silveria, S., Votavova, P., Salafsky, J., Pande, V.S., Kovar, M., Bluestone, J.A., Garcia, K.C. Engineering a Single-Agent Cytokine/Antibody Fusion That Selectively Expands Regulatory T Cells for Autoimmune Disease Therapy. J Immunol. 201, 2094–2106 (2018).

16. Silva, D.A., Yu, S., Ulge, U.Y., Spangler, J.B., Jude, K.M., Labão-Almeida, C., Ali, L.R., Quijano-Rubio, A., Ruterbusch, M., Leung, I., Biary, T., Crowley, S.J., Marcos, E., Walkey, C.D., Weitzner, B.D., Pardo-Avila, F., Castellanos, J., Carter, L., Stewart, L., Riddell, S.R., Pepper, M., Bernardes, G.J.L., Dougan, M., Garcia, K.C., Baker, D. De novo design of potent and selective mimics of IL-2 and IL-15. Nature. 565, 186–191 (2019).

17. Park, Y.H., Koo, S.K., Kim, Y., Kim, H.M., Joe, I.Y., Park, C.S., Kim, S.C., Han, D.J., Lim, D.G. Effect of in vitro expanded CD4(+)CD25(+)Foxp3(+) regulatory T cell therapy combined with lymphodepletion in murine skin allo-transplantation. Clin. Immunol. 135, 43–54 (2010).

18. Webster, K.E., Walters, S., Kohler, R.E., Mrkvan, T., Boyman, O., Surh, C.D., Grey, S.T., Sprent, J. In vivo expansion of T reg cells with IL-2-mAb complexes, induction of resistance to EAE and long-term acceptance of islet allografts without immunosuppression. J. Exp. Med. 206, 751–760 (2009).

19. Grinberg-Bleyer, Y., Baeyens, A., You, S., Elhage, R., Fourcade, G., Gregoire, S., Cagnard, N., Carpentier, W., Tang, Q., Bluestone, J., Chatenoud, L., Klatzmann, D., Salomon, B.L., Piaggio, E. IL-2 reverses established type 1 diabetes in NOD mice by a local effect on pancreatic regulatory T cells. J. Exp. Med. 207, 1871–1878 (2010).

20. Tang, Q., Adams, J.Y., Penaranda, C., Melli, K., Piaggio, E., Sgouroudis, E., Piccirillo, C.A., Salomon, B.L., Bluestone, J.A. Central role of defective interleukin-2 production in the triggering of islet autoimmune destruction. Immunity 28, 687–697 (2008).

21. Ruschak, A.M., Kay, L.E. Methyl groups as probes of supra-molecular structure, dynamics and function. J Biomol NMR 46, 75–87 (2010).

22. Sprangers, R., Kay, L.E. Quantitative dynamics and binding studies of the 20S proteasome by NMR. Nature 445, 618–622 (2007).

23. Ollerenshaw, J. E., Tugarinov, V., Kay, L. E. Methyl TROSY, explanation and experimental verification. Magn. Reson. Chem. 41, 843–852 (2003).

24. Gans, P., Hamelin, O., Sounier, R., Ayala, I., Durá, M.A., Amero, C.D., Noirclerc-Savoye, M., Franzetti, B., Plevin, M.J., Boisbouvier, J. Stereospecific isotopic labeling of methyl groups for NMR spectroscopic studies of high-molecular-weight proteins. Angew Chem Int Ed Engl. 49, 1958–62 (2010).

25. Vallurupalli, P., Sekhar, A., Yuwen, T., Kay, L.E. Probing conformational dynamics in biomolecules via chemical exchange saturation transfer: A primer. J Biomol NMR 67, 243–271 (2017).

26. Lundström, P., Vallurupalli, P., Religa, T.L., Dahlquist, F.W., Kay, L.E. A single-quantum methyl ^13^C-relaxation dispersion experiment with improved sensitivity. J Biomol NMR. 38, 79–88 (2007).

27. Sekhar, A., Kay, L.E. NMR paves the way for atomic level descriptions of sparsely populated, transiently formed biomolecular conformers. Proc Natl Acad Sci USA 110, 12867–74 (2013).

28. Henzler-Wildman, K.A., Thai, V., Lei, M., Ott, M., Wolf-Watz, M., Fenn, T., Pozharski, E., Wilson, M.A., Petsko, G.A., Karplus, M., Hübner, C.G., Kern, D. Intrinsic motions along an enzymatic reaction trajectory. Nature 450, 838−844 (2007).

29. Boehr, D.D., McElheny, D., Dyson, H.J., Wright, P.E. The dynamic energy landscape of dihydrofolate reductase catalysis. Science 313, 1638−1642 (2006).

30. Whittier, S.K., Hengge, A.C., Loria, J.P. Conformational motions regulate phosphoryl transfer in related protein tyrosine phosphatases. Science 341, 899−903 (2013).

31. Hammes, G.G., Chang, Y.C., Oas, T.G. Conformational selection or induced fit, a flux description of reaction mechanism. Proc. Natl Acad. Sci. USA. 106, 13737–13741 (2009).

32. Vogt, A.D., Di Cera, E. Conformational selection or induced fit? a critical appraisal of the kinetic mechanism. Biochemistry 51, 5894–5902 (2012).

33. Tilley, J.W., Chen, L., Fry, D.C., Emerson, S.D., Powers, G.D., Biondi, D., Varnell, T., Trilles, R., Guthrie, R., Mennona, F., Kaplan, G. LeMahieu, R.A., Carson, M., Han, R.-J., Liu, C.-M., Palermo R. Ju G. Identification of a small molecule inhibitor of the IL-2/IL-2Rα receptor interaction which binds to IL-2. J. Am. Chem. Soc. 119, 7589–7590 (1997).

34. Tugarinov, V., Kanelis, V., Kay, L.E. Isotope labeling strategies for the study of high-molecular-weight proteins by solution NMR spectroscopy. Nat. Protoc. 1, 749–754 (2006).

35. Natarajan, K., McShan, A.C., Jiang, J., Kumirov, V.K., Wang, R., Zhao, H., Schuck, P., Tilahun, M.E., Boyd, L.F., Ying, J., Bax, A., Margulies, D.H., Sgourakis, N.G. An allosteric site in the T-cell receptor Cβ domain plays a critical signalling role. Nat Commun. 8, 15260 (2017).

36. Sattler, M., Schleucher, J. Heteronuclear multidimensional NMR experiments for the structure determination of proteins in solution employing pulsed field gradients. Progress in Nuclear Magnetic Resonance Spectroscopy 34, 93–158 (1999).

37. Tugarinov, V., Kay, L.E. Ile, Leu, and Val methyl assignments of the 723-residue malate synthase G using a new labeling strategy and novel NMR methods. J. Am. Chem. Soc. 125, 13868–13878 (2003).

38. Rossi, P., Xia, Y., Khanra, N., Veglia, G., Kalodimos, C.G. ^15^N and ^13^C-SOFAST-HMQC editing enhances 3D-NOESY sensitivity in highly deuterated, selectively [^1^H,^13^C]-labeled proteins. J. Biomol. NMR 66, 259-271 (2016).

39. Delaglio, F., Grzesiek, S., Vuister, G.W., Zhu, G., Pfeifer, J., Bax, A. NMRPipe: a multidimensional spectral processing system based on UNIX pipes. J. Biomol. NMR 6, 277–293 (1995).

40. Vranken, W.F., Boucher, W., Stevens, T.J., Fogh, R.H., Pajon, A., Llinas, M., Ulrich, E.L., Markley, J.L., Ionides, J., Laue, E.D. The CCPN data model for NMR spectroscopy, development of a software pipeline. Proteins 59, 687–696 (2005).

41. Wang, X., Rickert, M., Garcia, K.C. Structure of the quaternary complex of interleukin-2 with its alpha, beta, and gammac receptors. Science 310, 1159–1163 (2005).

42. Waudby, C.A., Ramos, A., Cabrita, L.D., Christodoulou, J. Two-dimensional NMR lineshape analysis. Sci. Rep. 6, 24826 (2016).

43. Hansen, D.F., Vallurupalli, P., Lundstrom, P., Neudecker, P., Kay, L.E. Probing chemical shifts of invisible states of proteins with relaxation dispersion NMR spectroscopy: how well can we do? J. Am. Chem. Soc. 130, 2667–2675 (2008).

44. McDonald, L.R., Boyer, J.A., Lee, A.L. Segmental motions, not a two-state concerted switch, underlie allostery in CheY. Structure 20, 1363–1373 (2012).

45. Bouvignies, G., Kay, L.E. Measurement of proton chemical shifts in invisible states of slowly exchanging protein systems by chemical exchange saturation transfer. J Phys Chem B 116, 14311–14317 (2012).

46. Guenneugues, M., Berthault, P., Desvaux, H.A. method for determining B1 field inhomogeneity. Are the biases assumed in heteronuclear relaxation experiments usually underestimated? J Magn Reson 136, 118–126 (1999).

47. Bouvignies, G., Kay, L.E. A 2D ^13^C-CEST experiment for studying slowly exchanging protein systems using methyl probes: an application to protein folding. J Biomol NMR 53, 303–310 (2012).

